# Structural Reorganization and Relaxation Dynamics of Axially Stressed Chromosomes

**DOI:** 10.1101/2022.09.03.506488

**Authors:** Benjamin S. Ruben, Sumitabha Brahmachari, Vinícius G. Contessoto, Ryan R. Cheng, Antonio B. Oliveira Junior, Michele Di Pierro, José N. Onuchic

## Abstract

Micromechanical studies of mitotic chromosomes have revealed them to be remarkably extensible objects and informed early models of mitotic chromosome organization. We use a data-driven, coarsegrained polymer modeling approach, capable of generating ensembles of chromosome structures that are quantitatively consistent with experiments, to explore the relationship between the spatial organization of individual chromosomes and their emergent mechanical properties. In particular, we investigate the mechanical properties of our model chromosomes by axially stretching them. Simulated stretching led to a linear force-extension curve for small strain, with mitotic chromosomes behaving about ten-fold stiffer than interphase chromosomes. Studying the relaxation dynamics we found that chromosomes are viscoelastic solids, with a highly liquid-like, viscous behavior in interphase that becomes solid-like in mitosis. This emergent mechanical stiffness in our model originates from lengthwise compaction, an effective potential capturing the activity of loop-extruding SMC complexes. Chromosomes denature under large strains via unraveling, which is characterized by opening of large-scale folding patterns. By quantifying the effect of mechanical perturbations on the chromosome’s structural features, our model provides a nuanced understanding of *in vivo* mechanics of chromosomes.

## I. INTRODUCTION

Understanding the physical nature of chromosomes is an outstanding challenge in modern biology. The mechanical properties of chromosomes are relevant to their biological function, for example, during mitosis, when chromosomes are pulled and manipulated by spindle fibers, mechanical robustness to stretching is essential to preserve chromosome integrity. Recent experiments have probed the mechanical response of individual chromatin loci subjected to an externally applied force[1], revealing that interphase chromatin is highly dynamic and liquid-like. Other experiments probing chromatin mobility have given evidence that chromatin is arranged in a solid-like gel state[2]. It has further been shown that intact chromatin is needed to maintain the structural integrity of the whole nucleus[3] and that the mechanical responsiveness of the nucleus is modulated through heterochromatin formation [4].

The mechanical properties of mitotic chromosomes have been probed directly in experiments which use micromanipulation to stretch individual chromosomes and measure their force-extension curves [5–10]. Experiments have demonstrated that mitotic chromosomes extracted from live cells are highly extensible objects, maintaining their shape and native elasticity after being stretched up to five times their native length. Mitotic chromosomes display a linear force response in accordance with Hooke’s law for extensions up to 40 times their native length, followed by a force plateau [5]. Toward understanding the origin of chromosomal elasticity, several studies have explored how treatment of mitotic chromosomes with cleavage enzymes affects their mechanical properties [7, 11]. It was found that the elasticity of mitotic chromosomes was softened when treated with protease[11], which cleaves peptide bonds. However, the chromatin material disintegrated when treated with Micrococcal nuclease (MNase) [7], which cleaves DNA. These findings suggested that the DNA molecule itself was principally responsible for the structural integrity and observed mechanical properties of a chromosome rather than any contiguous protein scaffold [7]. For mitotic chromosomes assembled *in vivo*, measurements of the chromosome’s bending modulus agree with the expected bending modulus for a linear elastic rod with the same elastic modulus and radius [7], suggesting that the mitotic chromosome’s stiffness is distributed homogeneously over its cross-section. Recent technological advances have allowed for measurement of human chromosome mechanics with high force-resolution, revealing a nonlinear stiffening behavior in the low strain regime [10].

These initial investigations of their mechanical properties led to the “chromatin mesh” model for mitotic chromosomes [5], which describes the mitotic chromosome as a mesh of extensible chromatin fibers cross-linked by a variety of factors, like the SMC proteins. However, a growing list of experimental results has since demonstrated that the mitotic chromosome should not be treated as a homogeneous material; in particular, Ref. [12] suggests that mitotic chromosome stretching occurs between rigid “condensin centers” placed discontinuously along the chromosome’s length and Ref. [10] shows that TOP2A, another chromatin cross-linking protein, is not uniformly distributed along the mitotic chromosome’s length, leading to non-uniform stretching. In addition, recent super-resolution microscopy experiments have demonstrated that mitotic chromosomes are organized into ∼ 80 kb domains which remain intact when chromosomes are stretched to even 30 times their native length [13] These findings suggest that the chromatin mesh model’s explanation of chromosome elasticity is incomplete. It has been pointed out that at least part of the chromosomal elasticity may arise from the opening of large-scale chromatin folding [14], and hierarchical folding [10].

While early micro-mechanical studies have offered some insights into the internal organization of chromosomes, a revolution in conformation-capture technology has since produced a wealth of new information about chromosome organization [15–20]. In particular, Hi-C generates “contact maps” of the genome, which report the frequency with which any pair of genomic loci are observed to be spatially proximal. This technique allowed the discovery of rich structural motifs in the organization of the interphase nucleus, such as compartments and loops [15, 17, 21]. Hi-C-based studies suggest that chromosome organization in the interphase nucleus is intricately connected to the regulation of gene expression [22]. More recently, Hi-C has been applied to synchronous cell lines throughout mitotic cell division, to find that chromosome organization changes drastically when chromosomes are compacted into rod-shaped mitotic chromosomes [23, 24]. In particular, TADs and chromatin compartments disappear while SMC complex-driven chromatin looping (leading to lengthwise compaction) increases.

We previously introduced the Minimal Chromatin Model (MiChroM) [25], a data-driven, coarse-grained polymer model for chromosomes which uses the principle of maximum entropy to infer an energy landscape for chromosome folding from Hi-C contact maps. MiChroM has been shown to quantitatively recapitulate the rich patterns of contact observed in Hi-C experiments. Structural ensembles generated by MiChroM provided new insights into interphase chromosome organization through the interplay between phase separation of biochemically distinct segments of chromatin (compartmentalization) and the genomic distance-dependent lengthwise compaction attributed to the activity of motors such as condensin and cohesin (called the Ideal Chromosome Potential) [25–30]. MiChroM has been shown to be consistent with locus-locus distance distributions determined with fluorescence *in situ* hybridization (FISH) [25, 31] and the dynamical viscoelasticity of chromosomes observed in experiment [32]. Recent work has also shown MiChroM structures to be consistent with the structures observed in DNA-tracing experiments [33–35].

In this work, we study the relationship between a chromosome’s large-scale organization and its mechanical properties using computer simulations of chicken (DT40) chromosomes in interphase and in mitosis which are consistent with the Hi-C maps [23]. First, we test the mechanical properties of the chromosome with *in silico* measurements of its force-extension curve and bending rigidity. We observe a linear elastic response in the low-extension regime, with a spring constant that increases 10 fold from interphase to mitosis (fig. 1). The simulated chromosomes are found to unravel when chromosome extension reaches roughly double its native length, causing a plateau in their force-extension curves. The measured persistence length of 2.4µ*m* agrees with that expected for a homogeneous material, suggesting that stress is distributed evenly over the mitotic chromosome’s cross-section. Second, we investigate the dynamical properties of model chromosomes in a simulated stress-relaxation experiment. Chromosomes respond to force as a viscoelastic spring, with relaxation times in good agreement with experimental measurements on the order of seconds [6]. Finally, we investigate how stretching alters the chromosome’s internal organization. In the low-strain regime, we find that stretching perturbs large-scale chromosome organization while leaving local chromatin structure largely unaffected. Contact maps respond very differently to applied force in interphase and mitosis, suggesting distinct mechanisms of chromosome elasticity. Our work suggests the forces underlying the spatial organization of the chromosome are also responsible for their mechanical behavior.

**FIG. 1.**
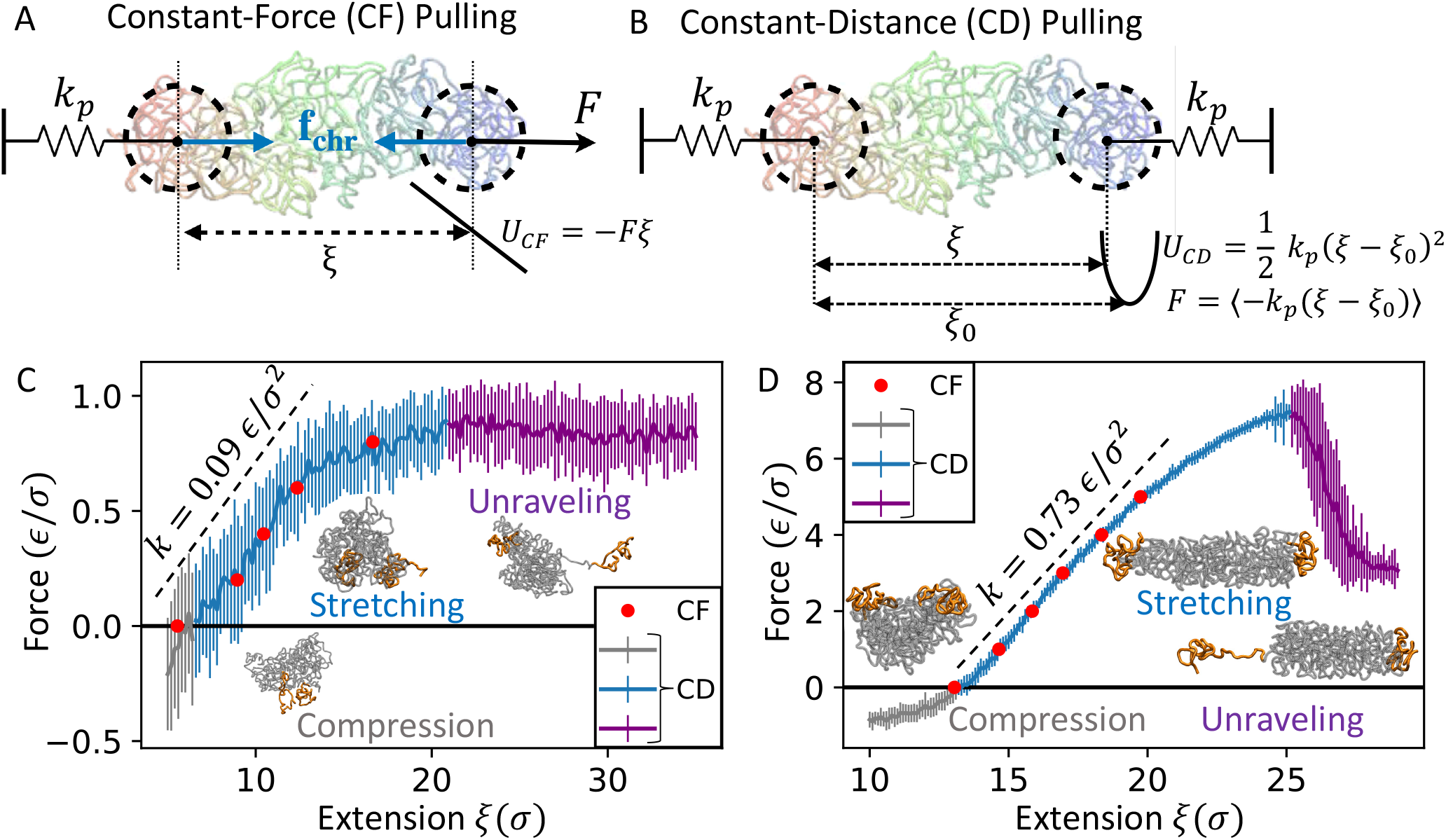
Force-Extension Behavior of Simulated Interphase and Mitotic Chromosomes. (A) Schematic representation of constant-force (CF) simulation setup. The reaction coordinate for stretching is defined as the x-distance between the centers of mass of the pull groups (*ξ* = *x*_*r*_ − *x*_*l*_). The pull groups are defined as the first and last 50 beads (2500 kb) of the chromosome. The center of mass of the left pull group (first 50 beads) is confined to the origin using a strong harmonic potential, and the right is constrained to the x-axis but allowed to slide along it. Constant is applied through a linear pulling potential *U*_*CF*_ = *Fξ* (B) Schematic representation of constant-distance (CD) simulations. Setup is the same as CF simulations, except chromosome extension is restrained to small window with a strong harmonic potential 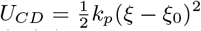. Force is measured as the average restoring force of the applied restraining potential (*F* = ⟨−*k*_*p*_(*ξ* − *ξ*_0_)⟩). (C/D) Resulting Force-extension curves for Interphase/Mitotic CF and CD measurements are overlaid. Force-extension curve for CD measurement is shown in grey, blue, and purple corresponding to compression, stretching, and unraveling regimes, respectively. Error bars show the standard deviation of the mean force over 40 replica ensembles. Red circles show force-extension curve for constant-force measurements. The initial slope of the force-extension curve is shown in black. Representative simulation snapshots for each regime are overlaid. Pull groups, defined as the first 50 and last 50 beads, are highlighted in orange.

## II. METHODS

Using the MiChroM energy function, we simulated DT40 chromosome 7 in both interphase and mitosis. Our model chromosomes are coarse-grained polymers consisting of 739 beads at 50 kb resolution. Chromosomes are simulated using Langevin dynamics at an effective temperature, subjected to a Hamiltonian made up of three components. The homopolymer potential includes bonding between neighboring polymers, bending rigidity of the polymer chain, and excluded volume interactions. Overlap between non-nearest neighbors is permitted with a finite energy penalty, allowing for topological relaxation of the polymer through chain-crossing. The type-type potential encodes interactions that depend on the epigenetic character of the participating segments of chromatin, resulting in phase-separation of chromatin into compartments. The ideal chromosome term encodes genomic-distance-dependent interactions which lead to lengthwise compaction of the chromosome, capturing the effect of loop-extruding motors such as cohesin and condensins [30]. We use parameters for the model that best agree with the experimental Hi-C maps of chicken DT40 cells (Fig. S1, S2). The simulations were carried out using the OpenMiChroM software package[36, 37]. OpenMiChroM is a Python library for performing chromatin dynamics simulations using GPU hardware acceleration through OpenMM Python API[38].

In order to probe the mechanical properties of these model chromosomes, we subjected them to axial stretching by applying pulling forces and orientation constraints to the centers of mass of the first 50 and last 50 beads of the polymer chain. Concretely, we write the centers of mass of the “pull groups” as follows:

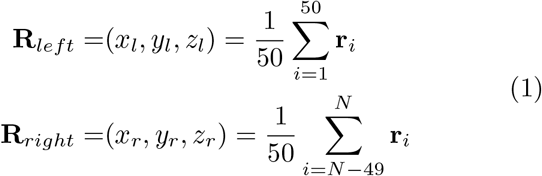

We define a pull coordinate *ξ* = *x*_*r*_ − *x*_*l*_, as the chromosome’s end-to-end x-distance. In chromosome pulling experiments, the ends of the chromosome are held by micropipettes and one is moved along a linear track. To mimic this process, we introduce orientation constraints:

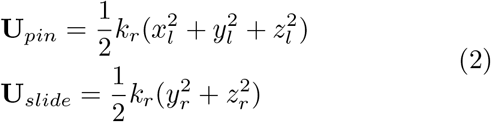

**U**_*pin*_ restrains the center of mass of the left pull group to the origin. **U**_*slide*_ restrains the right pull group to the x-axis, but allows it to slide along the x-axis. A large value *k*_*r*_ = 10^5^*ϵ/σ*^2^ is used so that **R**_*left*_ ≈ 0 and *y*_*r*_, *z*_*r*_ ≈ 0. We use two methods to subject the simulated chromosomes to axial strain. In constant-force (CF) pulling, a linear potential is used to apply a constant elongating force to the pull coordinate *ξ*. In constant-distance (CD) pulling, a harmonic potential with a very strong spring constant is used to constrain *ξ* to a very small window around a chosen reference distance *ξ*_0_.

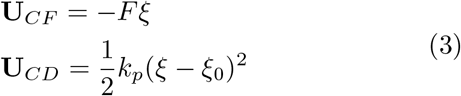

A large value of *k*_*p*_ = 10^5^*ϵ/σ*^2^ is chosen so that *ξ* ≈ *ξ*_0_ during CD sampling. These two pulling strategies are shown schematically in Fig. 1. These potentials acting on the centroids of the pull groups are implemented using the CustomCentroidBondForce class built into OpenMM [38]

In a CF simulation, chromosomes are equilibrated under the combined action of the MiChroM potential, **U**_*pin*_, **U**_*slide*_, and **U**_*CF*_. **U**_*CF*_ acts effectively to bias the pull coordinate *ξ* toward larger values, but does not constrain *ξ* to any particular value. An ensemble of equilibrium structures are recorded. During these simulations, *ξ* fluctuates over a relatively large interval. The histograms in figures S3A,D show the probability distributions *p*_*F*_ (*ξ*) of *ξ* during CF pulling at each chosen force value F. In order to create a force-extension curve, a single value *ξ*(*F*) must be chosen for each force F. The correct choice is *ξ*(*F*) = argmax [*p*_*F*_(*ξ*)], as explained in SI section VIII). In a CD simulation, chromosomes are equilibrated under the combined action of the MiChroM potential, **U**_*pin*_, **U**_*slide*_, and **U**_*CD*_. **U**_*CD*_ constrains the pull coordinate *ξ* to a very small window around a chosen reference distance *ξ*_0_ by exerting a restoring force *F*_*CF*_ = −*k*_*p*_(*ξ* − *ξ*_0_) on the pull coordinate. An ensemble of equilibrium structures are recorded, and the force is determined by averaging this restoring force over the generated ensemble:

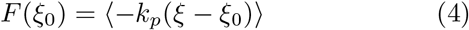

This force measurement method is justified more rigorously in SI section VII and figure S6.

## III. RESULTS AND DISCUSSION

### A. Chromosomes behave as linear elastic material in the small-strain regime

We measured the force-extension behavior of our model chromosomes using both the Constant-Force (CF) and Constant-Distance (CD) pulling methods (see methods). The resulting force-extension curves are shown in Figure 1 C-D. When subjected to low extensile axial force, model chromosomes behave like a linear elastic material. This means chromosome strain, defined as the ratio of the increase in the end-to-end extension to the native contour length, is linearly proportional to the applied axial force. The spring constant associated with the weak stretching of the chromosomes is obtained from the slope of the force-extension curve, with dimensions of force per unit length. We obtained spring constants of *k* = 0.09*ϵ/σ*^2^ and *k* = 0.73*ϵ/σ*^2^, respectively, for models of interphase and mitotic chromosomes, suggesting a near ten-fold stiffening of chromosomes in mitosis. Linearity of force-extension curves (Hooke’s law) has been experimentally observed for mitotic chromosomes [5, 14]. Similarly, recent rheology measurements in the interphase genome have indicated a weaker mechanical response [1]. The energy functions describing interphase and mitotic chromosomes are functionally identical. Parameters describing the properties of the chromatin chain (bond stiffness, bending rigidity, and excluded volume interactions) are identical across models, and the type-type and ideal chromosome potentials for interphase and mitosis diverge only in the numerical values of parameters describing the strength of compartmentalization and lengthwise compaction interactions (see Fig. S1). These parameters are learned solely from chromosome-contacts data, so that MiChroM recapitulates well the organization of interphase and mitotic chromosomes. The marked increase in stiffness from interphase to mitosis demonstrates that the chromosome’s mechanical properties are heavily influenced by its largescale organization.

The force required to stretch a mitotic chromosome to double its native length *In vitro* is typically in the range of 0.1 − 1 nN [14], which can be compared to the extrapolated doubling force of 9.5*ϵ/σ* for the simulated mitotic chromosome, suggesting that the reduced unit of force *ϵ/σ* ∼ 10 − 10^2^ pN. In SI section VI, we calibrate the model’s length scale (in mitosis) to be *σ* = 0.096µ*m* by matching the densities of the simulated and experimental mitotic chromosome. The reduced energy unit *ϵ* is an information-theoretic temperature which sets the energy scale of interactions between chromosome beads (containing hundreds of nucleosomes); in other words, the typical energy required to strain the interface between two 50 kb segments of chromatin is of the order of *ϵ*. Substituting *σ* = 0.096µ*m* into the relation *ϵ/σ* ∼ 10 − 10^2^ pN, we find *ϵ* ∼ 10^2^ − 10^3^*K*_*B*_*T* (recall *K*_*B*_*T* = 4.11 pN· nm).

Note that the CD method of pulling chromosomes provided a direct measure of the chromosome’s native length (Fig. 1). Whenever the two pull groups were closer than the native length of the chromosomes there was a compressive load on the external springs holding the pull groups. Simply put, in CD ensemble, the extension corresponding to the zero force in the force-extension curve gives the chromosome’s native length. We find that the native length of mitotic chromosomes is about 2-fold higher than that of interphase chromosomes.

### B. Chromosomes unravel under strong axial stretching

When stretched to lengths near double their native length, model chromosomes lose their structural integrity as the polymer separates into blobs connected by a single-bead-thick chromatin thread. In interphase, the unraveling leads to a plateau in the force-extension curve. In mitosis, there appears a sharp dip in the force-extension curve at the onset of unraveling, followed by a plateau (See figure 1). Chromatids assembled *in vitro* from Xenopus egg extracts have been observed to segregate into domains of thick chromatin connected by thin filaments when stretched to lengths greater than 15 times their native length [8]. A blob-and-thread geometry was also observed in stretched mitotic newt chromosomes when treated with MNase (an enzyme that cuts DNA) [14]. For model chromosomes, this rupture can be understood as a consequence of free-energy minimization. As the chromosome stretches uniformly, its surface area increases, decreasing the number of (energetically favorable) bead-to-bead contacts. When stretched past rupture, the chromosome segregates into a “dense” phase, where the chromosome contacts are preserved, and a “rare” phase characterized by a bare segment of polymer. Rupture occurs when the energetic benefit of preserving native contacts in the dense phase outweighs the entropic cost of phase separation (see Fig. S14 E). The sudden drop in the restoring force upon unraveling of mitotic chromosomes suggests that the transition to the coexistence of dense and rare phases is abrupt. For the interphase chromosome, the onset of phase coexistence is smooth, likely due to its lower mechanical stiffness.

### C. Stress relaxation in simulated chromosomes is well captured by a linear viscoelastic model

We measured the dynamics of chromosome extension in a stress-relaxation simulation, and found that the chromosome response to sudden stress is well captured by a linear-viscoelastic Kelvin-Voight (KV) model (Fig. 2). In this model, a linear dashpot with viscosity *η* and a linear (Hookean) spring with spring constant *k* are attached in parallel. When the KV model is subjected to an external stress *F*, the end-to-end extension *ξ* evolves in time following: 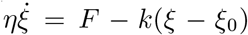. After the sudden application of stress to the chromosome, its extension *ξ* increases at an exponentially decaying rate : *ξ*(*t*) = *ξ*_0_ + (*F/k*) (1 − exp(−*t/τ*_*r*_)). The saturation extension, *ξ*(*t* → ∞) = *ξ*_0_ + *F/k*, controlled by the spring constant, increases linearly with the applied stress according to Hooke’s law. After the pulling force is suddenly released, the chromosome’s extension returns to its native length following an exponential decay: *ξ*(*t*) = *ξ*_0_ + (*F/k*) exp(−*t/τ*_*r*_). In Fig. 2, we fit the trajectories of the chromosome’s extension to these equations, and determine the relaxation time *τ*_*r*_, which are 1.2 sec. and 18 sec. respectively for mitosis and interphase. These relaxation times represent the time scale over which all the internal modes of fluctuations of the chromosomes relax, and the perturbed extension reaches equilibrium. Due to their higher mechanical stiffness, compact mitotic chromosomes relax faster than the less compact interphase chromosomes.

**FIG. 2.**
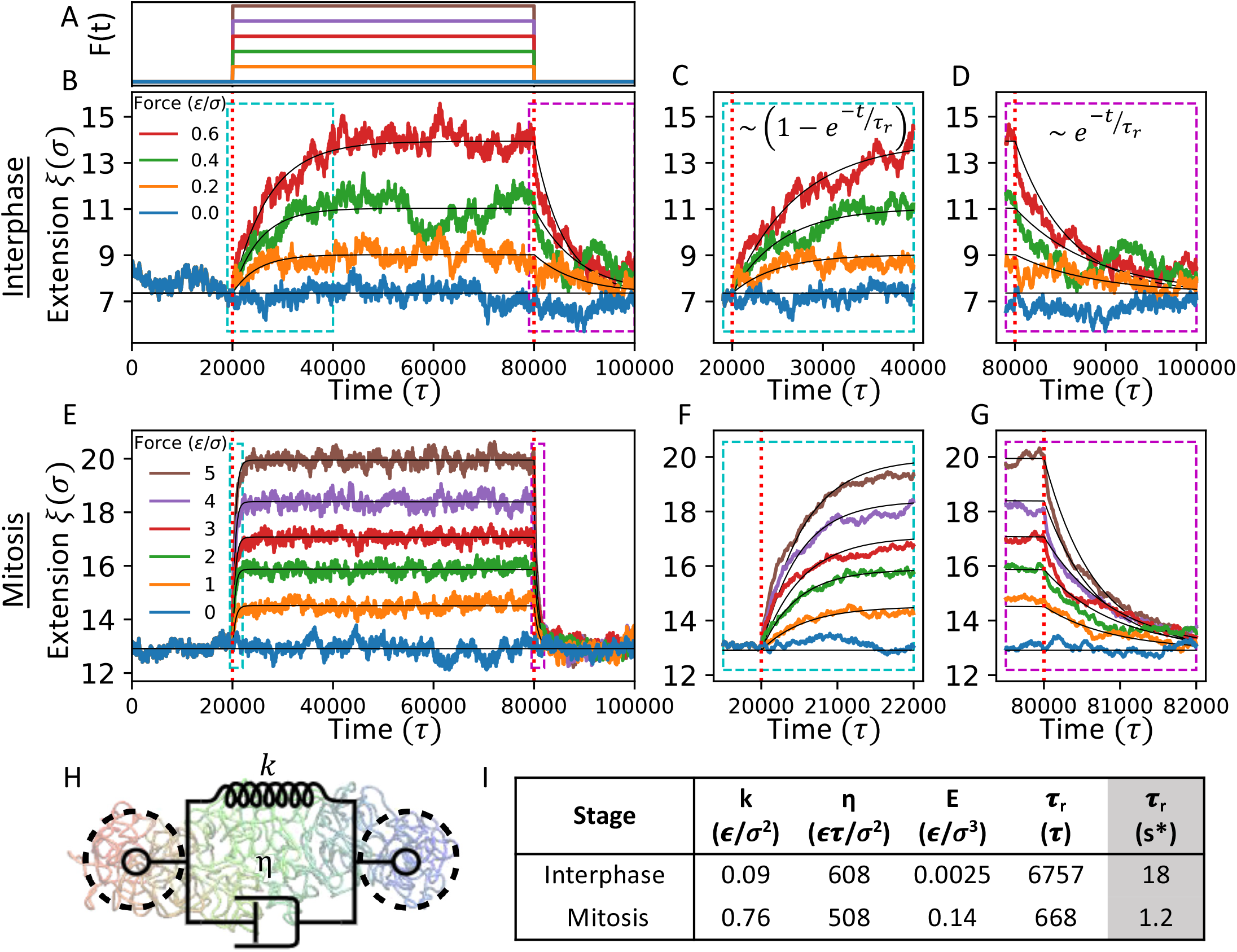
Viscoelastic Relaxation Dynamics of Simulated Chromosomes. (A) F(t) exerted on chromosomes during stress relaxation experiments. For interphase, force values are (in reduced units) 0, .2, .4, .6 (*F* = .8 not shown due to chromosome unraveling). For prometaphase, force values are (in reduced units) 0, 1, 2, 3, 4, 5. (B/E) Mean Extension vs. Time plot during stress-relaxation experiment for interphase. Force is applied at time 2000*τ* and released at 8000*τ*. Chromosome extension *ξ* shown is averaged over 40 replica trajectories to reduce noise. Thin black lines show fit of mean chromosome extensions to the exponential decays indicated in the expressions at the top right. (C/F) More detailed view of chromosome’s response to the sudden force onset. (D/G) More detailed view of chromosome’s response to the sudden force release. (H) Chromosome Behavior is compared to that of a simple Kelvin-Voigt model with a spring constant and viscosity. (I) Kelvin-Voigt parameters of best fit. *k* is spring constant, *η* is viscous damping constant. The intrinsic analogs of these parameters (elastic modulus E and relaxation time*τ*_*r*_) are listed as well. Quantities are expressed in reduced units. *Relaxation time is also converted to units of seconds (See SI). For mitotic chromosome, measured relaxation time of 1.2*s* matches well with experimental results. Note that in figure 2, the trajectories *ξ*(*t*) shown are averaged over 40 replica trajectories of each model chromosome. The trajectories of individual chromosomes are compared to the mean in figure S5.

We note that the relaxation times measured in simulation are in good agreement with those estimated for a free-draining polymer. When a tension *F* is applied to the chromosome of spring constant *k*, its extension will increase by a distance of *d* = *F/k*. Thus, the chromosome’s center of mass will change by a distance of *d*_*mean*_ = *F/*2*k*. If relaxation occurs with a characteristic timescale *τ*_*r*_, the mean center-of-mass velocity during relaxation is approximately *v*_*cm*_ ≈ *d*_*mean*_*/τ*_*r*_ = *F/*(2*kτ*_*r*_). Chromosome 7 has 739 beads each of mass 10, and with friction constant of *γ* = 0.1*τ*^−1^. The net drag of the chromosome’s center of mass is *ζ* = *Nmγ*. Therefore the total drag force on the chromosome during stretching is approximately *F*_*drag*_ ≈ *ζv*_*cm*_. Setting the drag force and applied force equal, we obtain a relaxation time estimate of *τ*_*r*_ ≈ *Nmγ/*2*k*. The reduced time unit *τ* is calibrated in SI section VI, note that the conversion factor to real time units is larger in interphase than mitosis, due to the different length scales of the models. Experiments probing the mitotic chromosomes find a relaxation timescale of roughly 2 seconds [6], which is in excellent agreement with the value obtained from simulation (Fig. 2I).

### D. Stretching Perturbs Interphase Chromosomes at the Site of Pulling, while Mitotic Chromosomes Distribute Stress along their Axial Length

Having quantified the mechanical properties of our model chromosomes, we now seek to understand the mechanisms which underlie their elasticity. To do so, we use a variety of structural analyses to observe how the chromosome’s internal structure is perturbed by axial stretching. In Figure 3, we plot Hi-C contact maps of interphase and mitotic chromosomes in the absence of applied tension, with an applied tension, and the residual difference map. In the absence of applied tension, the interphase chromosome’s telomeres co-localize under the combined influence of compartmentalization interactions favoring contacts between loci of like epigenetic character and the entropic favorability of end-to-end proximity induced by their confinement to the x-axis (see SI section IX). When tension is applied, the telomeres move apart leading to a depletion of contact probability at the anti-diagonal extremities of the contact map (Fig. 3 A-C). In contrast, the mitotic chromosome distributes applied tension along its length, leading to a nearly translation-invariant pattern of contact probability changes in its Hi-C contact map (Fig. 3 E-G). Contacts are depleted at all genomic separations except for a band around 3-5 Mb of separation where tension *increases* contact probability (see Fig. S4 bottom right). Stretched mitotic chromosomes have a straighter backbone (Fig. 4) which may enhance structural helicity, leading to a more prominent non-monotonic scaling of contact probability. Supplemental videos 1 and 2 show how interphase and mitotic chromosome contact maps change as chromosomes are stretched using the CD (constant-distance) pulling method.

**FIG. 3.**
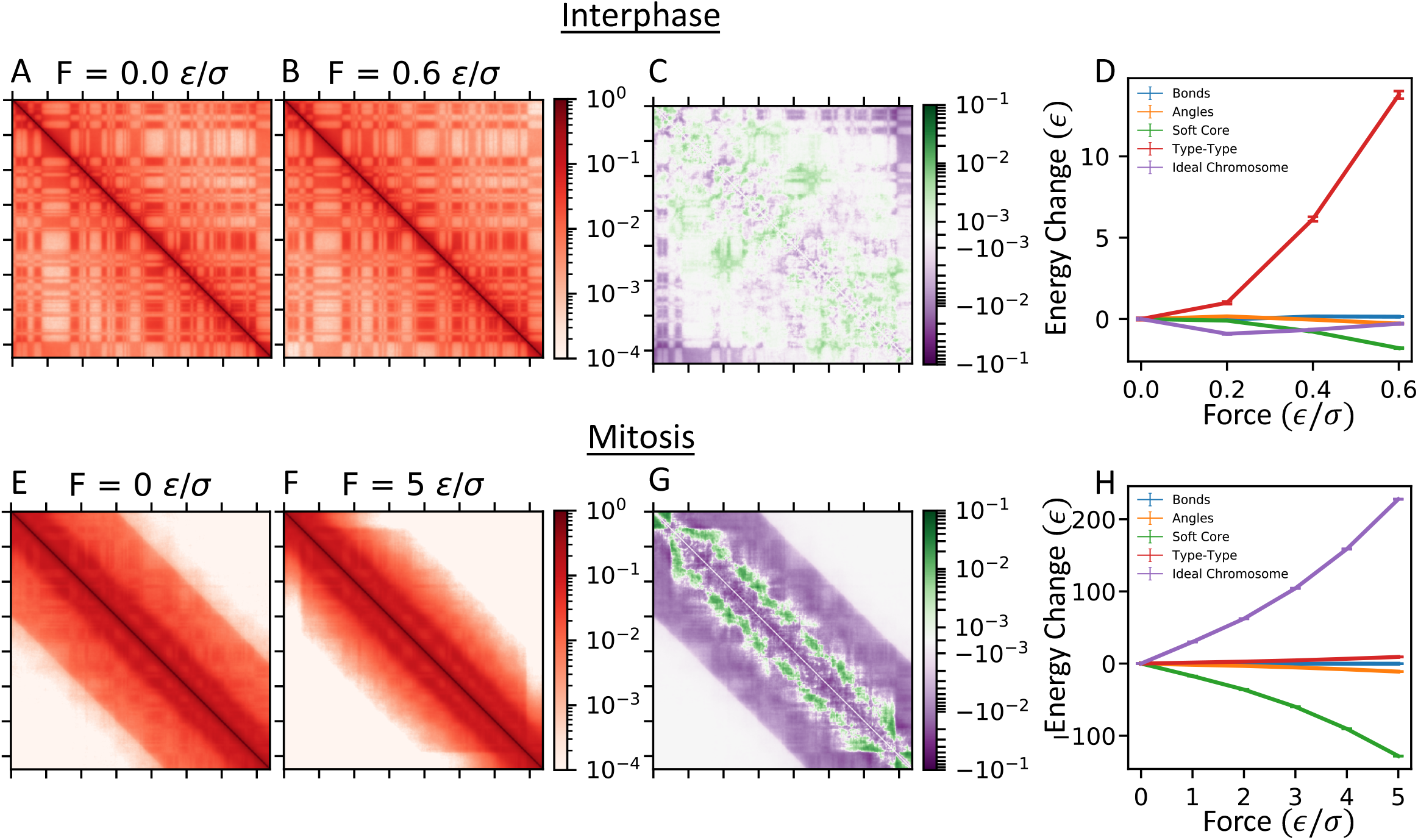
Perturbations to Chromosome’s contact map and potential energy components under tension for interphase (top) and mitosis (bottom).(A/E) Simulated Hi-C Contact maps of the interphase/prometaphase chromosome under CF setup with applied force 0. (B/F) Simulated Hi-C Contact maps of the interphase/prometaphase chromosome under CF setup with applied force 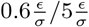. (C/G) The difference between the contact maps for chromosomes under tension (B/F) and not under tension (A/E). Contact probability plotted on a symmetric log scale with linear cutoff 10^−3^. (B/E) Change in the interphase/prometaphase chromosome’s mean contact probability vs. genomic distance curve under tension relative to the structure under no tension. Vertical dotted line shows genomic distance beyond which all contacts are between the two groups. In interphase, pulling mainly disrupts interactions between the two pull groups. In prometaphase, pulling has almost no effect on the pull groups’ interactions, but leads to a distinct pattern of lost contacts around 200 kb and 10000 kb as well as a slight enhancement of interactions around 4000 kb. (C/F) Mean potential energy components of the interphase/prometaphase chromosome under tension. Bonds, Angles, and Soft-Core potential energy components are included in the homopolymer potential. In interphase, pulling primarily disrupts Type-Type interactions which depend on the epigenetic types of interacting loci. In prometaphase, pulling primarily disrupts Ideal-Chromosome interactions which depend on the genomic distance between loci.

**FIG. 4.**
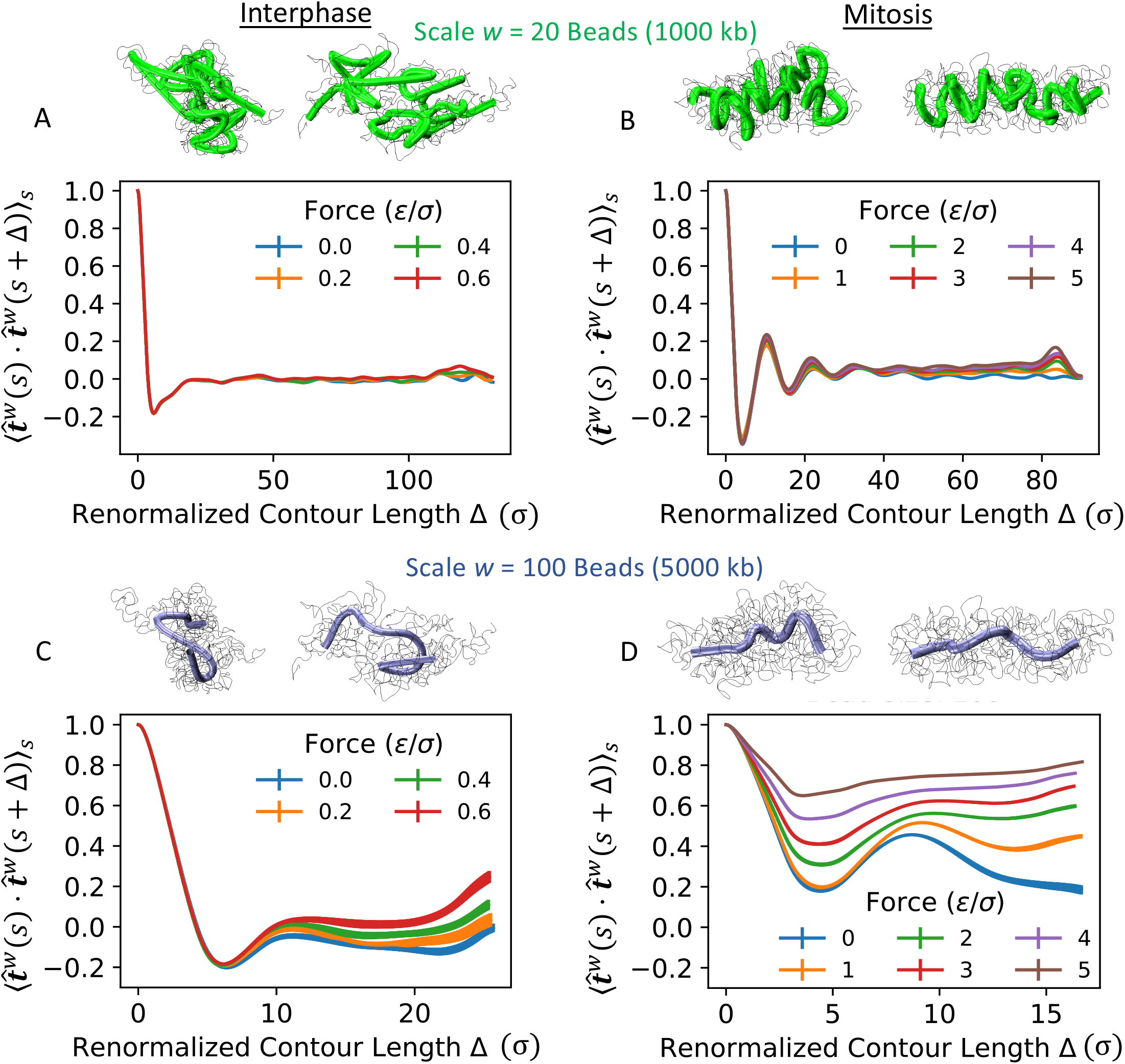
Pulling causes reorganization of large-scale chromosome architecture. (A/C) Effect of tension on the interphase chromosome renormalized at the scale of *w* = 20 / *w* = 100 beads (1000 kb / 5000 kb). Above plots, representative simulation snapshots are shown under force *F* = 0 (left) and 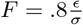 (right). Full chain is shown as a thin black line with renormalized chain 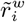 shown as a tube. Plots show renormalized tangent correlation functions 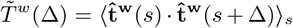 for chromosomes under constant force *F* = 0, 0.2, 0.4, 0.6. (B/D) Effect of tension on the mitotic chromosome renormalized at the scale of *w* = 20 / *w* = 100 beads (1000 kb / 5000 kb). Above plots, representative simulation snapshots are shown under force *F* = 0 (left) and 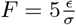 (right). Full chain is shown as a thin black line with renormalized chain 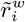 shown as a tube. Plots show renormalized tangent correlation functions 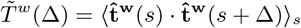 for chromosomes under constant force *F* = 0, 1, 2, 3, 4, 5.

### E. Compartmental Interactions Mediate Interphase Chromosome Elasticity while Lengthwise Compaction Drives Mitotic Chromosome Elasticity

We also examined the effects of stretching on each component of the MiChroM energy functional. In Figure 3C/3F, we plot the interphase and mitotic model’s mean potential energy components as a function of the externally applied tension, relative to the zero-tension ensembles. Pulling interphase chromosomes results mainly in an increase in the type-type potential energy component whereas pulling mitotic chromosomes results mainly in an increase in the ideal-chromosome component. This suggests that distinct mechanisms underlie the elasticity of interphase and mitotic chromosomes: interphase chromosome elasticity arises from mechanical rigidity of chromatin compartments while mitotic chromosome elasticity arises from SMC-driven lengthwise compaction. Note that a decrease in the average soft-core potential, which represents self-avoidance between non-neighboring monomers, is a direct result of the depletion of inter-monomer contacts in stretched chromosomes.

### F. Chromosomes Undergo Large-Scale Rearrangements under Tension

To further investigate the relationship between chromosome elasticity and structure, we examine the structural rearrangements at different length scales in a stretched chromosome. We introduce the “renormalized” chain, which is a lower resolution version of our original chain obtained by averaging the polymer bead positions over a window of *w* beads. The renormalized chain at scales *w* = 20 beads (1000 kb) and *w* = 100 beads (5000 kb) are visualized in Fig. 4.

We analyzed the bond correlation function at different coarse-graining lengths: 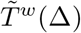 (Fig. 4). This quantity, bound between -1 and 1, is the mean cosine of the angle made by a pair of bonds that are Δ distance apart along the contour. We expect a decay for neighbors because of the stiffness of the polymer, while the non-monotonic behavior after the initial decay is due to looping back of the chromosome. Note that the non-monotonic oscillations depict a long-range helical order along the chromosome backbone, which is enhanced in mitosis. With increasing stretching force, the bonds align along the external force, but the reorientation is significant only at lengthscales above 100 beads (∼5 Mb) in interphase and 50 beads (∼2.5 Mb) in Mitosis, suggesting significant perturbation of the structure only at these large length scales. Our analysis of the linear compaction along the renormalized chain *C*(*w*), defined as the ratio of contour length of the full polymer to the contour length of the renormalized chain at scale *w*, and the radius of gyration of the renormalized window, are consistent with the 100/50 bead (∼5/2.5 Mb) lengthscale as a characteristic length for structural perturbation under small strain in interphase/mitosis (See SI section I, Fig. S7-S13).

### G. Mitotic Chromosomes Behave as Stiff Elastic Rods

Bending rigidity and persistence lengths are two important coefficients that characterize the mechanical response of chromosomes [5, 8, 14]. We measured the persistence length (*L*_*p*_) and bending rigidity (*B*) of our model mitotic chromosomes (recall, *B* = *K*_*B*_*TL*_*p*_ [39]). To allow for accurate measurement of persistence length, we extended the length of the chromosome by joining 10 copies in series (and allowing for type-type and ideal chromosome interactions between concatenated chromosomes, effectively creating a single elongated chromosome). Note that, as persistence length is an intensive property, increasing the chromosome’s length will not affect the persistence length. The persistence length of chromosomes that are reported experimentally is related to the bending persistence of the chromosome axial backbone. Hence, we calculated the core or backbone of the mitotic chromosome by averaging the bead positions over a sliding window of *w* = 560 beads. This window size was chosen so that the diameter of the chromosome equals the mean renormalized bond length (See SI section I D and Fig. S15 for details). We then calculate the persistence length by fitting the bond vector correlations obtained at the coarse-graining lengthscale of *w* = 560 to the following equation: 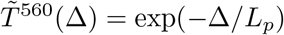.

Our analysis reports the persistence length associated with backbone bending of the mitotic chromosome to be (*L*_*p*_)_*measured*_ = 21.1*σ* ≈ 2.0*µm* (Fig. 5B). This is in excellent agreement with the expected persistence length of a linear material with given radius (*r* = 3.24*σ*) and force constant (*f*_0_ = 9.5*ϵ/σ*): (*L*_*p*_)_*expected*_ = *f*_0_*r*^2^*/*(4*K*_*B*_*T*) = 25*σ* ≈ 2.4*µm*. This suggests that as the mitotic chromosome bends, stress is distributed homogeneously over its cross-section. Experimental measurements reached the same conclusion for mitotic chromosomes extracted from Newt and Xenopus cells [14, 40]. The persistence length of DT40 mitotic chromosomes has not been measured, but a persistence length of 2*µm* appears visually consistent with images of mitotic chromosomes 15m after release from G2 arrest (see Fig. 1 in [23]). The obtained persistence length of 21.1*σ* is approximately 3.3 times the diameter of the mitotic chromosome. This is much smaller than the ratio measured for mitotic chromosomes extracted from Newt and Xenopus cells [14, 40], but matches well the experimentally measured ratio of 3.4 for chromatids assembled from Xenopus egg extracts [8]. Zhang et. al use a similar strategy to measure the persistence length of another model for the mitotic chromosome, similarly finding persistence lengths from 1.5-4 times the diameter of their model chromosomes [41]. Metaphase chromosomes retrieved from living Xenopus or newt cells had much larger persistence lengths ranging from millimeters to centimeters [7]. The lower values of persistence length in our prometaphase model may also be attributed to partially condensed chromosomes in prometaphase compared to metaphase.

**FIG. 5.**
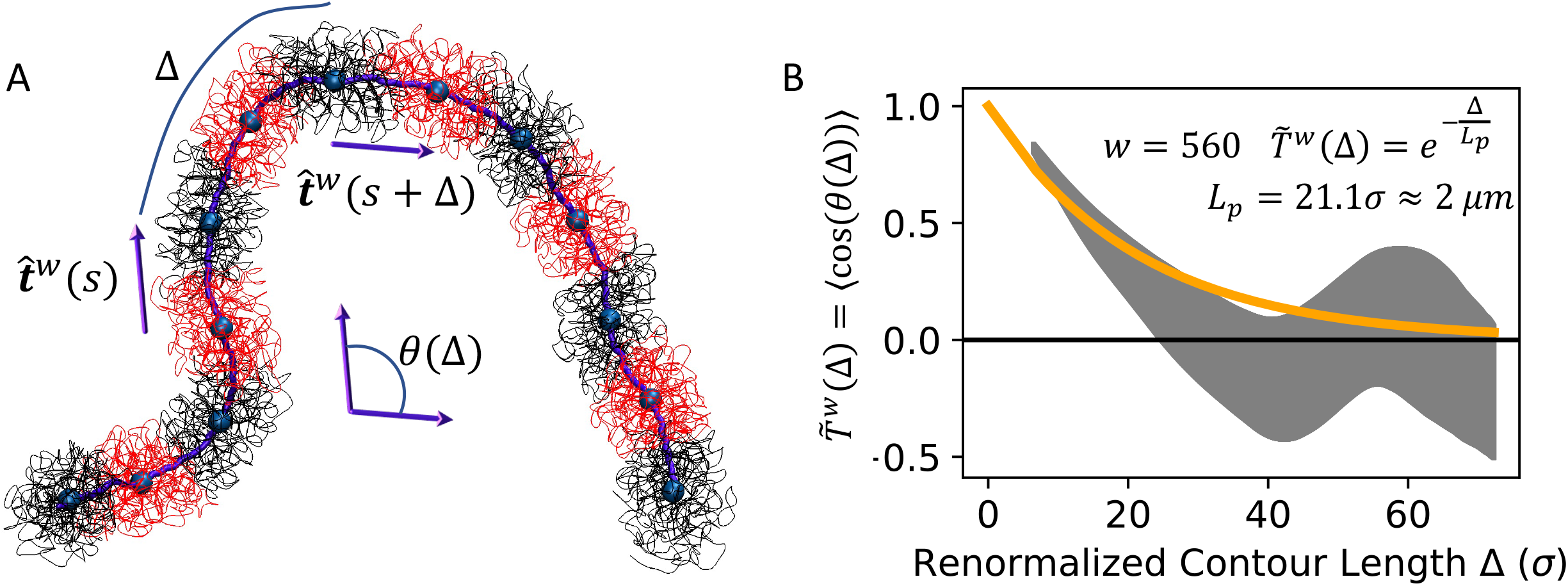
Bending rigidity of the prometaphase chromosome. (A) Simulation Snapshot of 7390-bead mitotic chromosome. Renormalized chain 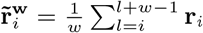 for *l* ∈ {1, 2, …, *N* − *w* + 1} is shown in purple. One set of nearest non-overlapping neighbor renormalized beads 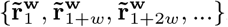 is shown as large blue spheres. Blue spheres are the centers of mass of the associated chromatin windows shown in alternating red and black thin tubing. A natural scale of *w* = 560 beads is chosen self-consistently so that chromosome radius equals mean renormalized bond length (the distance between blue spheres, see fig S15 for details). Purple arrows represent renormalized bonds used to calculate the renormalized tangent correlation function. (B) Renormalized tangent correlation function of the mitotic chromosome’s thin core, calculated according to equation 13. Gray region shows standard error of the mean over four replica trajectories. Correlation function is fit to a decaying exponential to determine persistence length.

## IV. DISCUSSION

In this work, we have tested the mechanical properties of a bead-spring polymer model of chromosomes with data-driven force fields [25] by subjecting it to external stretching, mimicking tweezers-style experiments [5, 14]. The data-driven force field comprises of three components: first, the homopolymer potential contains terms such as the nearest neighbor bonds and inter-monomer self-avoidance; second, the phase separation potential that drives compartmental segregation; and third, lengthwise compaction, encoded by the ideal chromosome potential, that crumples the polymer resembling the steady-state loop-extrusion activity of SMC complexes[30]. Note that the prometaphase chromosomes of chicken DT40 cells show a non-monotonic trend in the scaling of contact probability with genomic distance, suggesting an increase in contact frequency between genomic segments that are ∼ 4 Mb apart [23]. We find that using a non-monotonic ideal chromosome potential is necessary to recapitulate the high intensity stripe pattern running parallel to the diagonal (see Fig. S1).

We pull the chromosomes using two different techniques: constant force (CF) and constant distance (CD), and show that our results are consistent (Fig. 1). Both interphase and mitotic chromosomes show a linear force-extension curve in the small-strain regime, behaving like a linear elastic material. However, mitotic chromosomes are about ten-fold stiffer than the interphase ones under stretching perturbations (Fig. 1). The native contour length, i.e., the end-to-end distance for zero stretching force, of mitotic chromosomes is about 2 fold higher than interphase chromosomes. Matching the mitotic chromosome’s extrapolated doubling force measured in simulations to the typical doubling force observed in experiments, we back-solve to estimate the reduced energy unit, finding *ϵ* ∼ 10^2^ − 10^3^*K*_*B*_*T*. This value sets the energy scale for interactions between beads representing 50 kb of chromatin. The density of crosslinking proteins in mitotic chromosomes has been estimated as approximately 1 connector protein every 6 kb of chromatin [5, 42]. Thus we expect the pairwise interactions between beads in MiChroM to capture the action of approximately 10 cross-linking proteins. An estimate of *ϵ* ∼ 10^2^ − 10^3^*K*_*B*_*T* is reasonable in this context.

When stretched to more than twice their native contour length (i.e., larger than 100% strain), the chromosomes begin to unravel into a blob-and-thread structure. The unraveling force for mitotic chromosomes is about tenfold higher than interphase. Similar unraveling has been observed in experiments, though at very high strains [5, 14]. Mechanical perturbations of interphase chromosomes are much less explored experimentally, nonetheless, initial studies indeed suggest a mechanical rigidity that is much weaker than mitotic chromosomes [1].

When subjected to a sudden stress, model chromosomes show slow relaxation just like a visco-elastic material (Fig. 2). Using a linear-viscoelastic Kelvin-Voight model, we were able to extract relaxation times corresponding to interphase (∼ 20 sec) and mitotic (∼ 2 secs) chromosomes (Fig. 2). The faster relaxation of mitotic chromosomes may be attributed to their higher mechanical stiffness that leads to a faster decay of the fluctuation modes. The extracted relaxation times for mitotic chromosomes are in good quantitative agreement with experimental findings [14].

Stretching perturbs the structure of interphase and mitotic chromosomes in qualitatively distinct ways. While the weak restoring force upon stretching interphase chromosomes arises from disrupting compartments and separating the telomeres; the stiff response from stretching mitotic chromosomes comes from SMC-driven lengthwise compaction (Fig. 3). Interestingly, our model predicts that the stripe pattern in the mitotic Hi-C map is enhanced upon stretching (Fig. 3). The stripe, controlled to condensin II activity [23], signifies emergence of helical order along the backbone of the chromosomes. When there is no stretching force, the entropy of the backbone obscures its helical order, however, as the entropic excursions of the backbone are suppressed by external force, the signature corresponding to the helical order becomes stronger. This is a direct prediction of our model and remains to be experimentally validated.

Force-induced structural rearrangements in our model occur at lengthscales near 100 beads or 5 Mb (Fig. 4). This suggests that the source of linear elasticity in our model comes from rearrangements of large chunks of DNA that maintain their internal hierarchical organization, but displace relative to each other. We also measured the bending rigidity and persistence length of our model prometaphase chromosomes (Fig. 5). The measured values are consistent with the theoretical expectation from a linear elastic rod of the structure, however, experimental observations for chicken DT40 prometaphase chromosomes are lacking at the moment.

The reversible linear elasticity of mitotic chromosomes appears to span a wider strain regime in experiments [5] than our model. This may arise from the fact that MiChroM is trained to reproduce *population-averaged* Hi-C contact maps, so that mechanically robust contacts in single cells are averaged into weaker population-wide spatial associations. MiChroM treats all cell-cell variability in chromosome structure as *annealed* disorder. While we simulate many copies of a chromosome in parallel to speed sampling, in principle the population-averaged contact map could be recovered as the long-time average contact map of a single chromosome. The extent to which individual chromosomes explore the space of folded conformations *in vivo* is at present unclear. Single-cell interphase Hi-C contact maps have demonstrated large variability in chromosome structure between cells of the same population, and have much sparser contact maps than population-averaged Hi-C [43]. In addition to annealed disorder, cell-cell variability in epigenetic markings or random placement of stably binding crosslinkers may create *quenched* disorder between cells of the same population. If trained on these sparse single-cell Hi-C contact maps, an energy landscape model might learn a comparatively sparse set of strong interactions that resemble stable crosslinkers. Future work may investigate the mechanical properties of energy landscape models trained on single-cell Hi-C data in this manner.

Our simulations should be compared to the simulations of Zhang et. al [41], who developed a “dynamic loop model” for mitotic chromosomes. This model consists of a polymer chain on a lattice. When two loci come into close spatial proximity, a cross-linking interaction can form between them if they are separated by a contour length less than a chosen cutoff. This leads to cylindrical structures resembling mitotic chromosomes. Cross-linking interactions have a finite lifetime, and thus effectively resemble the transient interactions created by the ideal chromosome potential in MiChroM. Similarly to MiChroM, these model chromosomes display linear force-extension behavior at small strain, but unravel when strained past 3 times their native length, opening into a blob-and-thread geometry. The stretch elasticity of dynamic loop chromosomes depended sensitively on the concentration of crosslinkers. Also like the MiChroM mitotic chromosome, dynamic loop chromosomes have a persistence length that is only a few times their diameter.

Experiments have demonstrated that condensin compacts chromatin by actively extruding loops, deliberately bringing together loop anchors much more quickly than passive diffusion [44]. However, both MiChroM [25] and the dynamic loop model [41] rely on passive diffusion to bring loop anchors into close spatial proximity. Chromosome models which include active extrusion have been implemented to explain TAD formation in interphase [45, 46] and the compaction and separation of sister chromatids during mitosis [47]. The mechanical properties of this class of models remain unexplored.

## V. CONCLUSION

In our simulations, interphase and mitotic chromosomes are both modeled as bead-spring polymers with additional pairwise short-range interactions tuned to recreate Hi-C contact maps. Despite differing only in the numerical values of interaction parameters, simulated interphase and mitotic chromosomes appear to stretch through distinct mechanisms. The interphase chromosome responds to pulling mainly at the site of stretching, as the compartments formed by the telomeres are torn apart. The mitotic chromosome, by contrast, stretches through the depletion of both genomically distant and genomically adjacent contacts throughout its length, while contacts around 4000 kb remain intact. Up to nearly 2x extension, large-scale rearrangements allow the mitotic chromosome to stretch its end-to-end distance without losing its structural integrity.

The Minimal Chromatin Model is inferred from Hi-C contact maps, and trained only to generate an ensemble of structures consistent with experimentally measured structures. *A priori*, there is little reason to expect that MiChroM structures that satisfy contact patterns also properly describe the chromosome’s mechanical properties. Yet, MiChroM mitotic chromosomes recapitulate many nontrivial mechanical properties of mitotic chromosomes: they reproduce the shape of the mitotic chromosome’s force-extension curve for nearly two-fold extensions, display slow viscoelastic relaxation dynamics, and have stretch and bending moduli related as those of an ideal elastic rod.

Stretched chromosome structures differ from native structures through rearrangements in large-scale chromosome architecture. The qualitative agreement between theory and experiment in the low-extension regime raises the possibility that mitotic chromosomes stretch through large-scale rearrangements of chromosome architecture *in vivo*. Indeed, spindle-generated forces are known to stretch mitotic chromosomes to only double their native length in large animal and insect cells [9], a regime well captured by MiChroM.

Other interesting observations can be gleaned by comparing our results for interphase and prometaphase chromosomes: we observe a 50-fold increase in the chromosome’s elastic modulus upon condensation from interphase to mitosis, and the interphase and prometaphase force-extension curves have distinct shapes. These differences illuminate a connection between chromosome organization and chromosome micromechanics: the mechanisms which drive chromosome organization at the micro-scale are disrupted during chromosome stretching, leading to an elastic response. As interphase chromosomes are primarily organized through type-type compartmentalization and mitotic chromosomes are primarily organized through SMC-driven lengthwise compaction, their responses to pulling forces vary drastically. In particular, the “bump” in the mitotic chromosome’s contact scaling curve around 4000 kb creates an anchor of enhanced interactions which maintain the mitotic chromosome’s structural integrity during stretching.

In all, we studied the varied mechanical response of chromosomes as they are organized into interphase- or mitosis-specific structures. We quantify the emergent stiffness in mitotic chromosomes, and attribute it to the SMC-driven lengthwise compaction. We also found that chromosomes behave as viscoelastic solids, as captured by a Kelvin-Voight model, where the interphase chromosomes are more liquid-like or viscous than their mitotic counterparts. Our study quantifies the mechanical aspects of the MiChroM model that will help guide future modeling efforts. Furthermore, our estimates for forces and time scales underlying features like compartments is important for a nuanced understanding of the in vivo mechanics of chromosomes.

## Supporting information

Supplemental Video 1: Simulated Hi-C in Stretched Interphase Chromosome

Supplemental Video 2: Simulated Hi-C in Stretched Mitotic Chromosome

## ACKNOWLEDGMENTS

This work was supported by the Center for Theoretical Biological Physics sponsored by the National Science Foundation NSF NSF (Grants PHY-2019745 and CHE-1614101) and by the Welch Foundation (Grant C-1792). JNO is a Cancer Prevention and Research Institute of Texas (CPRIT) Scholar in Cancer Research. ABOJ acknowledges the Robert A. Welch Postdoctoral Fellow program. BSR was supported in part by the Rice University department of physics and astronomy’s Summer Undergraduate Research Fellowship, as well as the Rice University Center for Theoretical Biological Physics’ Opportunities for Research in Biophysics, Informatics, and Theoretical Science (ORBITS) summer program. BSR was also supported by the National Institutes of Health Molecular Biophysics Training Grant NIH/ NIGMS T32 GM008313.

