## Supplemental Information for "Structural Reorganization and Relaxation Dynamics of Axially Stressed Chromosomes"

(Dated: August 28, 2022)

### I. STRUCTURAL ANALYSIS SPANNING LENGTH SCALES

In this section, we describe a strategy for analyzing polymer structure at length scales from the resolution of the model up to arbitrarily large-scale features. Throughout this section, $\langle\cdot\rangle_i$ is used to refer to averages over an index $i$ which runs over monomer indices and $|\cdot|$ refers to the euclidean norm of a vector.

#### A. Renormalized Chains

The configuration of a polymer is specified by the N 3-D coordinates of its monomers. Suppose these positions are given by $\mathbf{r}_i$ for $i = 1, ...N$. To study the properties of this chain at different length scales, our strategy will be to divide the chain into segments of $w$ beads, and take the average positions of these segments:

$$[1/w\left(\mathbf{r}_1 + \cdots + \mathbf{r}_w\right), 1/w\left(\mathbf{r}_{w+1} + \cdots + \mathbf{r}_{2w}\right), \dots] \quad (1)$$

To obtain more measurements at the scale of $w$, we can also shift our frame of reference for dividing the chain:

$$\begin{aligned}&[1/w\left(\mathbf{r}_2 + \cdots + \mathbf{r}_{w+1}\right), 1/w\left(\mathbf{r}_{w+2} + \cdots + \mathbf{r}_{2w+1}\right), \dots];\\ &[1/w\left(\mathbf{r}_3 + \cdots + \mathbf{r}_{w+2}\right), 1/w\left(\mathbf{r}_{w+3} + \cdots + \mathbf{r}_{2w+2}\right), \dots]\\ &\dots\end{aligned} \quad (2)$$

and so on. To incorporate information from each frame of reference, we define $\tilde{\mathbf{r}}^{\mathbf{w}}$, the "renormalized chain at scale $w$" by averaging the positions of the polymer's monomers over a *sliding* window of w beads:

$$\tilde{\mathbf{r}}^{\mathbf{w}}_i = \frac{1}{w}\sum_{l=i}^{i+w-1}\mathbf{r}_l \quad \text{for} \quad i \in \{1, 2, ..., N-w+1\} \quad (3)$$

We define the renormalized bonds at scale w as:

$$\tilde{\mathbf{v}}^{\mathbf{w}}_i = \tilde{\mathbf{r}}^{\mathbf{w}}_{i+w} - \tilde{\mathbf{r}}^{\mathbf{w}}_i \quad (4)$$

and the corresponding unit tangent vectors as:

$$\hat{\mathbf{t}}^{\mathbf{w}}_i = \frac{\tilde{\mathbf{v}}^{\mathbf{w}}_i}{|\tilde{\mathbf{v}}^{\mathbf{w}}_i|} \quad (5)$$

Note that the $i+w$ subscript in equation 4 ensures that renormalized bonds go between the centers of mass of adjacent, non-overlapping windows.

For each window of $w$ beads, we can calculate a radius of gyration:

$$[R_g]_i(w) = \left(\frac{1}{w}\sum_{l=i}^{i+w-1}(\mathbf{r}_i - \tilde{\mathbf{r}}^{\mathbf{w}}_i)^2\right)^{\frac{1}{2}}$$

Averaging over the index i gives a mean radius of gyration for all windows of width w:

$$R_g(w) = \langle [R_g]_i(w)\rangle_i$$

#### B. Renormalized Bond Lengths and Compaction

Between any two bead indices $i$ and $j$, we can associate a renormalized contour length between the two indices:

$$\tilde{L}^w_{ij} = \sum_{k=i}^{j-1}\frac{1}{w} * |\tilde{\mathbf{v}}^{\mathbf{w}}_k| \quad (6)$$

Averaging this quantity over all pairs $\langle l, l+\delta\rangle$ with an index separation $\delta$, we obtain

$$\tilde{L}^w_\delta = \langle \tilde{L}^w_{l,l+\delta}\rangle_l \approx \frac{\delta}{w} * \langle|\tilde{\mathbf{v}}^{\mathbf{w}}_l|\rangle_l \quad (7)$$

*
†

which can be thought of as the average contour length at scale $w$ between beads of index separation $\delta$. Here, we assumed that the mean renormalized bond lengths at scale $w$ are approximately equal at all positions along the chain.

We define the chromosome's linear compaction at scale $w$ as:

$$C(w) = \frac{w * \tilde{L}_1^1}{\tilde{L}_w^w} \tag{8}$$

Equivalently, the linear compaction at scale $w$ is the ratio of the chromosome's contour length to the contour length of the renormalized chain at scale $w$.

#### C. Renormalized Tangent Correlations

Now, between any two bead indices $i$ and $j$ we can associate a renormalized tangent correlation, defined through the following equation:

$$\tilde{T}_{ij}^w = \hat{\mathbf{t}}_i^{\mathbf{w}} \cdot \hat{\mathbf{t}}_j^{\mathbf{w}} \tag{9}$$

Averaging over all pairs $\langle l, l+\delta \rangle$ with an index separation $\delta$, we obtain

$$\tilde{T}_\delta^w = \langle \hat{\mathbf{t}}_l^{\mathbf{w}} \cdot \hat{\mathbf{t}}_{l+\delta}^{\mathbf{w}} \rangle_l \tag{10}$$

In equations 10 and 7, we see that the index difference $\delta$ parameterizes a renormalized tangent correlation function:

$$\tilde{T}^w(\Delta) = \langle \hat{\mathbf{t}}^{\mathbf{w}}(s) \cdot \hat{\mathbf{t}}^{\mathbf{w}}(s+\Delta) \rangle_s \tag{11}$$

that depends on renormalized contour length $\Delta = \tilde{L}_\delta^w$. In this equation, $s$ represents the distance along the renormalized contour, which depends implicitly on the monomer index through equation 7. This is the function plotted in figures 4, 5 and S6.

It is well known that semiflexible chains in thermal equilibrium have an average tangent correlation function of the form:

$$\tilde{T}^w(\Delta) = \langle \hat{\mathbf{t}}^{\mathbf{w}}(s) \cdot \hat{\mathbf{t}}^{\mathbf{w}}(s+\Delta) \rangle_s = e^{-\Delta/l_p} \tag{12}$$

where $l_p$ is the persistence length of the semiflexible chain. We use this relationship to fit the persistence length of simulated mitotic chromosomes, which take the form of a long polymer folded into a flexible cylinder (see main text Fig. 5).

#### D. Renormalized Radius of a Bending Cylinder

For cylindrical chromosomes, we are presented with the challenge of calculating the radius of a bending cylinder. One way to do this is to first calculate a renormalized chain at a sufficiently large scale $w$ to serve as a central axis. We then measure the distances of individual beads from this central axis. The distance of the i'th bead from the central axis $\tilde{\mathbf{r}}^{\mathbf{w}}$ is given by:

$$r_i^w = \min_{j=1,\ldots,N-w} |\mathbf{r}_i - \tilde{\mathbf{r}}_j^{\mathbf{w}}| \tag{13}$$

We calculate radial distances $r_i^w$ for each bead i, and compile them into a histogram of distances from the central axis. From this histogram, we define the radius $R^w$ as the radial distance at which the cumulative distribution reaches .95. Precisely, $R^w$ is defined implicitly through the following relationship:

$$\mathbb{E}_{i\in\{1,\ldots,N\}}[\Theta(R^w - r_i^w)] = .95 \tag{14}$$

Here, $\Theta$ is the Heaviside step function. For an example, see figure S15. Note that the radius measurement depends on the chosen scale w. This method is used to calculate a radius of $R = 3.24\sigma$ for the extended mitotic chromosome (see main text Fig. 5). However, the mitotic DT40 chromosome 7 is too short to reliably calculate a thin core. In this case, we replace the thin core with the x-axis, as the orientation restraints align the short chromosome with the x-axis. This leads to a radius measurement of $R = 4.82\sigma$ for the original-length DT40 chromosome 7.

### II. REDUCED UNITS

Our simulations are performed using reduced units. The reduced unit of energy, $\epsilon$, is set to unity, and defines the thermal energy scale $K_BT$ of the system. As such, our generated ensembles of structures are sampled from a Boltzmann distribution with energy scale $\epsilon$ such that the probability of observing a state i with energy $E_i$ is given by: $P_i \propto e^{-E_i/\epsilon}$. Lengths are measured in reduced units $\sigma$. The diameters of the simulation beads (corresponding to the width of a single bead representing 50 kb of DNA) are set to $1\sigma$. Time is measured in reduced units $\tau$. All other units (e.g. mass) can be derived from these reduced units.

### III. SIMULATION DETAILS

We perform molecular dynamics simulations of DT40 chromosome 7 in both interphase (G2) and mitosis (prometaphase, 15 minutes after release from G2 arrest). The chromosome is modeled using the Minimal Chromatin Model, to be described shortly. The Minimal Chromatin model is a coarse-grained polymer model in which each simulation bead represents 50 kb of chromatin. For DT40 chromosome 7, this results in a 739-bead polymer. Molecular dynamics simulations were implemented with

Open-MiChroM, a fast and scalable platform for GPU-accelerated chromosome simulation described in Ref. [36]. Open-Michrom is based in the OpenMM python API [38] for molecular dynamics simulations.

The models for interphase and mitotic chromosomes were trained using Hi-C contact maps generated by Ref. [23]. The training procedure in Ref. [25] was used to infer an effective energy landscape for the 739-bead polymer that recreates Hi-C contact maps with high precision. Chromatin types annotations, were also obtained for this chromosome from [23]. The Gene Expression Omnibus (GEO) accession number for the data sets used in training is GSE102740.

#### A. The Minimal Chromatin Model (MiChroM)

The Minimal Chromatin Model Hamiltonian is given by:

$$U_{\mathrm{MiChroM}}\left(\vec{r}\right)=U_{HP}(\vec{r})+\sum_{\substack{k\geq l\\ k,l\in\text{ Types}}}\alpha_{kl}\sum_{\substack{i\in\{\text{ Loci of Type }k\}\\ j\in\{\text{ Loci of Type }l\}}}f\left(r_{ij}\right)+\chi\cdot\sum_{(i,j)\in\{\text{ Loops Sites }\}}f\left(r_{ij}\right)+\sum_{d=3}^{500}\gamma(d)\sum_{i}f\left(r_{i,i+d}\right) \quad (15)$$

with the contact function:

$$f\left(r_{ij}\right)=\frac{1}{2}\left(1+\tanh\left[\mu\left(r_c-r_{ij}\right)\right]\right) \quad (16)$$

In this work, we use parameters $\mu = 1.51$ and $r_c = 2.12$ and the set of chromatin types are simplified to two structural types A and B. The interaction parameters $\alpha_{kl}$ governing type-type interaction strengths are given by:

| | Interphase | Mitosis |
|---|---|---|
| $\alpha_{AA}$ | -0.1309 | -0.0164 |
| $\alpha_{AB}$ | -0.0789 | -0.0020 |
| $\alpha_{BB}$ | -0.1530 | -0.0151 |

with types assigned to the 739 beads as follows:

AAAAAAAAABABBBBBBBBBBBBBBBAABBBBBBBBBBBBBBBAAAAAAAAAAAAAAAABAA
BBBBBBBBBBBBBBBBBBBBBAAAAAAAAAAAAAAAAAAAAAAAAAAAAAAAAAAAAAAAAA
AAAAAAAAAAAAAAAAAAAAAAAAAAAAAAAAAAAAAAAAABBBBBBBBBBBBBBBBBBBBB
BBBBAAAABAAABBAAAAAAAAAABBBBBBBBBBAAAAAAAAAAAAAAAAAAAAAAAAAAAA
AAAAAABBBBBBBBBBBBBBBAAAAAAAAAABBBBAAAAAAAAAABBBBBBBBBBBBBBBBB
BBBBBAAABBBBBBBAAAABBBBBAAAAAAAAAAAAAAAAAAAAAAAAAAAAAAAABBBBBA
AAAABBBBABBBBBBBBBBBBBBBBBBBBBBBBBBBBBBBBBBBBBBBBBBBBBBBBBAABB
BAAABBBAAAAAAAAAAAAAAAAAABAABBBBBAAAAAAAAAAAABBBBBBBBBBBBBBBBBB
BBBBBBBBBBBAAAAAAABBBBBBBBBBAAAAAAAAAAAAAAAAAAAABAABBBABBBBBBB
BBAAAAAAAAAAAAABBBBBBAAABBBBBBBBBBBBBAABBBBBBBBBBAAABBAAAAAAAA
ABBBBBBBBBBABBBBBBBBBBBBBBBBBBBBBBBBAABBBBBBAAAAAAAABABBBBBBBB
BBBBBBBBBBBAAAAAAAAAAAAAAAAAAAAAAAAAAAAAAAAAAAAAAAAAAAAAAAAAAA

The ideal chromosome interaction parameters $\gamma(d)$ are plotted in figure S1.

#### B. The Homopolymer Model

The Homopolymer Model captures generic polymer behavior, and is written as follows:

$$\begin{aligned}U_{HP}(\vec{r})=&\sum_{i\in\{\text{ Loci}\}}U_{FENE}\left(r_{i,i+1}\right)+\sum_{i\in\{\text{ Loci}\}}U_{hc}\left(r_{i,i+1}\right)\\&+\sum_{i\in\{Angles\}}U_{\text{Angle}}\left(\theta_i\right)+\sum_{i,j\in\{\text{ Loci}\}}U_{sc}\left(r_{i,j}\right)\end{aligned} \quad (17)$$

The $U_{FENE}$ (Finitely Extensible Nonlinear Elastic) contribution is the bonding potential between neighboring monomers.

$$U_{FENE}\left(r_{i,j}\right)=\begin{cases}-\frac{1}{2}k_bR_0^2\ln\left[1-\left(\frac{r_{i,j}}{R_0}\right)^2\right] & \text{for } r_{i,j}\leq R_0\\ 0 & \text{for } r_{i,j}>R_0\end{cases} \tag{18}$$

Neighboring monomers also experience a purely repulsive hard-core excluded volume potential that prevents overlap between consecutive beads:

$$U_{hc}\left(r_{i,j}\right)=\begin{cases}4\varepsilon\left[\left(\frac{\sigma}{r_{i,j}}\right)^{12}-\left(\frac{\sigma}{r_{i,j}}\right)^6+\frac{1}{4}\right] & \text{for } r_{i,j}\leq\sigma 2^{\frac{1}{6}}\\ 0 \quad \text{for } r_{i,j}>\sigma 2^{\frac{1}{6}}\end{cases} \tag{19}$$

Polymer stiffness is added through a three-body potential that acts on all sets of three consecutive beads:

$$U_{\text{Angle}}\ \left(\theta_i\right)=k_a\left[1-\cos\left(\theta_i-\theta_0\right)\right] \tag{20}$$

Finally, non-bonded pairs of beads experience a soft-core repulsion that allows for occasional chain crossing, which is essential to capture the effect of topoisomerases:

$$U_{sc}\left(r_{i,j}\right)=\begin{cases}\frac{1}{2}E_{cut}\left[1+\tanh\left(\frac{2U_{LJ}(r_{i,j})}{E_{cut}}-1\right)\right] & r_{i,j}<r_0\\ U_{LJ}\left(r_{i,j}\right) & r_0\leq r_{i,j}\leq\sigma 2^{\frac{1}{6}}\\ 0 & r_{i,j}>\sigma 2^{\frac{1}{6}}\end{cases}$$
$$U_{LJ}\left(r_{i,j}\right)=4\varepsilon\left[\left(\frac{\sigma}{r_{i,j}}\right)^{12}-\left(\frac{\sigma}{r_{i,j}}\right)^6+\frac{1}{4}\right] \tag{21}$$

$r_0$ is set to the distance where $U_{LJ}(r_0)=\frac{1}{2}E_{\text{cut}}$.

In this work, we use the following parameter values:

$$k_a=\epsilon\ \ k_b=30\epsilon/\sigma^2\ \ E_{\text{cut}}=4\epsilon\ \ R_0=1.5\sigma \tag{22}$$

### IV. MOLECULAR DYNAMICS SIMULATIONS OF DT40 CHROMOSOME 7

Chromosomes are initialized from an extended spiral state (see figure S16). We first condense chromosomes by running a 50,000 step molecular dynamics simulation with the potential:

$$U_{\text{Condense}}\ (\vec{r})=U_{\text{MiChroM}}\ (\vec{r}) + \sum_{i=1}^{N}\frac{1}{2}k_s*(|\mathbf{r}_i|-r_0)^2\Theta(|\mathbf{r}_i|-r_0) \tag{23}$$

where $|\mathbf{r}_i|$ is the distance of bead i from the origin and $\Theta$ is a Heaviside step function. We choose a value of $k_s=0.2\frac{\epsilon}{\sigma^2}$ and the value of $r_0$ is chosen to fix the mean density inside the sphere of radius $r_0$ as 0.1 beads/$\sigma^3$.

To orient the chromosome along the x-axis, we then run a local energy minimization, followed by a 50,000 step molecular dynamics simulation at a temperature $k_BT=3\epsilon$ under the following potential:

$$U_{\text{Orient}}(\vec{r})=U_{\text{MiChroM}}(\vec{r})+U_{pin}(\mathbf{R}_{left}) + U_{slide}(\mathbf{R}_{right})-f_rx_r\Theta(x_0-x_r) \tag{24}$$

Here, $U_{pin}(\mathbf{R}_{left})$ and $U_{slide}(\mathbf{R}_{right})$, given in the main text, restrict the left pull group's center of mass to the origin and the right pull group's center of mass to the x-axis. The last term applies a constant force to the center of mass of the right pull group along the positive x-axis if its x-coordinate is below $x_0$. Values $f_r=100\epsilon/\sigma$ and $x_0=7\sigma$ are chosen. This potential ensures that the "right" pull group (last 50 beads) are oriented in the positive x-direction relative to the "left" pull group (first 50 beads).

Finally, to produce equilibrated initial structures for simulations, we run a 1,000,000 step simulation at temperature $k_BT=\epsilon$ under the following potential:

$$U_{\text{Equilibrate}}(\vec{r})=U_{\text{MiChroM}}(\vec{r})+U_{pin}(\mathbf{R}_{left})+U_{slide}(\mathbf{R}_{right}) \tag{25}$$

The starting structures obtained here are used as inputs for the remaining simulations. This process is repeated 40 times for interphase and 40 times for mitosis to give 40 different starting structures for simulations of each phase of the cell cycle. We term these starting structures "oriented structures."

#### A. Open Simulations

Open simulations are standard MiChroM simulations under the MiChroM potential with no added orientation restraints or pulling forces. (We term these "open" because we do not use a spherical hard-wall potential as has been done in previous works). Starting from the oriented structures, we simulate chromosomes for 60,000,000 steps under the standard MiChroM potential:

$$U_{\text{Open}}(\vec{r}) = U_{\text{MiChroM}}(\vec{r}) \tag{26}$$

After the first 6,000,000 steps (which allow the chromosome to equilibrate in the absence of $U_{pin}(\mathbf{R}_{left})$ and $U_{slide}(\mathbf{R}_{right})$), we collect structures every 3000 steps, to give a total of 18,000 structures per simulation. 40 replica trajectories are produced. The resulting in-silico Hi-C contact maps are compared to experimental contact maps in Fig. S1, and in the scatter plots in figure S2B/E.

#### B. Native Simulations

Native simulations are simulations under the MiChroM potential with added restraints that pin the left pull group to the origin and pin the right pull group to the x-axis, but allow the right pull group to slide along the x-axis. (We term these "native" because they can be used to measure the native length of the chromosome in the presence of orientation restraints).

Starting from the oriented structures, we simulate chromosomes for 5,000,000 steps under the following potential:

$$U_{\text{Native}}(\vec{r}) = U_{\text{MiChroM}}(\vec{r}) + U_{pin}(\mathbf{R}_{left}) + U_{slide}(\mathbf{R}_{right}) \tag{27}$$

We record the x-distance between pull groups $(x_r - x_l)$ every 5 steps and save chromosome conformations every 500 steps. 40 replica trajectories are produced. The resulting in-silico Hi-C contact maps are compared to experimental contact maps in figure S2A/D.

#### C. Stress Relaxation Simulations (Constant Force)

Stress relaxation simulations use the following potential:

$$U_{\text{Stress-Relaxation}}(\vec{r}, t) = U_{\text{MiChroM}}(\vec{r}) + U_{pin}(\mathbf{R}_{left}) + U_{slide}(\mathbf{R}_{right}) - F(t)(x_r - x_l) \tag{28}$$

$$F(t) = \begin{cases} 0 & \text{if } t < 20,000\tau \text{ or } t \geq 80,000\tau \\ F & \text{if } 20,000\tau \leq t < 80,000\tau. \end{cases} \tag{29}$$

the forces F used are (in reduced units) F = 0, .2, .4, .6, .8 for interphase and F = 0, 1, 2, 3, 4, 5 for mitosis. We run 40 replica trajectories of each chromosome. The F=0.8 ensemble in interphase is left out of some analysis due to unraveling of many replica chromosomes at this force (see figure S4). During simulations, the reaction coordinate is recorded every 100 steps ($1\tau$). Polymer conformations are saved every 1000 steps ($10\tau$) between time $35,000\tau$ and $80,000\tau$ to generate ensembles of structures under constant force (CF ensembles).

#### D. Constant-Distance Simulations

To generate ensembles of chromosome conformations stretched to constant extensions, we must first generate extended starting structures. To do so, we perform a constant-velocity pulling experiment. Starting from the previously equlibrated oriented structures, we run a local energy minimization and 300,000-step equilibration under constant potential $U_{\text{CV}}(\vec{r}, t = 0)$ (see equation below). Then, we run a constant-velocity pulling simulation under time-dependent potential:

$$U_{\text{CV}}(\vec{r}, t) = U_{\text{MiChroM}}(\vec{r}) + U_{pin}(\mathbf{R}_{left}) + U_{slide}(\mathbf{R}_{right}) + \frac{1}{2}k_p(x_r - x_l - \xi_0(t))^2 \tag{30}$$

$$\xi_0(t) = \xi_{\text{start}} + v * t \tag{31}$$

For interphase chromosomes, we use $\xi_{\text{start}} = 5\sigma$ and $v = .0002\sigma/\tau$ For prometaphase chromosomes, we use $\xi_{\text{start}} = 10\sigma$ and $v = .0002\sigma/\tau$. Conformations are saved every 50,000 steps, yielding structures with reaction coordinates distributed by distances of $0.1\sigma$. Velocities are chosen to be much smaller than the inverse relaxation timescales calculated in stress-relaxation experiments, so that chromosomes remain in (or near) quasi-static equilibrium during constant-velocity pulling. Ultimately, confirmation that chromosomes have equilibrated comes from the excellent agreement between CF and CD force-extension curves (see main text figure 1).

These extended structures are then used as initial conformations for simulations with constant reference distance (CD simulations). We simulate chromosomes for 5,000,000 steps under the following potential:

$$U_{\text{CD}}(\vec{r}) = U_{\text{MiChroM}}(\vec{r}) + U_{pin}(\mathbf{R}_{left}) + U_{slide}(\mathbf{R}_{right}) + \frac{1}{2}k_p(x_r - x_l - \xi_0)^2 \tag{32}$$

The reaction coordinate $\xi$ is recorded every 5 steps and polymer conformations are saved every 500 steps, giving 50,000 polymer conformations per simulation. Because we choose a very large vale of $k_p = 10,000\epsilon/\sigma^2$, $\xi = x_r - x_l \approx \xi_0$ throughout the simulation. At extension $\xi_0$, force is measured according to equation 4. 40 replica simulations are run at each reference distance. For the interphase chromosome, we chose 130 reference distances $\xi_0 = 5, 5.2, 5.4, ..., 35$. For the mitotic chromosome, we chose 190 reference distances $\xi_0 = 10, 10.1, 10.2, ..., 29$ (see figure S6A/B).

### V. EXTENDED MITOTIC CHROMOSOME SIMULATION TO CALCULATE PERSISTENCE LENGTH

Because the mitotic DT40 chromosome 7 is very short, it isn't possible to properly measure its persistence length. Therefore, we simulate an "extended" version of the mitotic chromosome by concatenating 10 copies of DT40 chromosome 7 into a chain 7390 beads long. On the extended mitotic chromosome, we perform only standard "open" simulations under potential $U_{\text{MiChroM}}(\vec{r})$. An initial structure is generated as a condensed spiral conformation (see figure S16). To destroy any bias that comes from the initial structure, we simulate for 9,000,000 steps at temperature $k_BT = 3\epsilon$, then for 9,000,000 steps at temperature $k_BT = 2\epsilon$. Finally, to equilibrate, we simulate for 60,000,000 steps at $k_BT = \epsilon$. During this last equilibration step, bead masses are reduced from 10 to 1 to speed the approach to equilibrium. Finally, using the equilibrated structures as initial configurations, we simulate for 500,000 steps, saving polymer conformations every 50,000 steps. 4 replica trajectories are produced.

### VI. REAL UNITS

Previously, FISH experiments have been used to calibrate the length scale in MiChroM for interphase chromosomes of the cell line GM06990. In this work, we simulate a different cell line and introduce a model for the prometaphase chromosome. Because the size of a single 50 bk bead is fixed at $\sigma$, but local chromatin density can change between cell types and cell cycle stages, prior length scale calibrations do not necessarily transfer.

We calibrate the length scale of the prometaphase chromosome by matching the density of the chromosome to the density of mitotic chromosomes observed in electron micrographs: 1 nucleosome (200 bp) per 11nm x 11nm x 11nm cube. We find that a value of $\sigma = 0.096\mu m$ leads to good agreement between simulated and experimental chromatin densities.

We can now calibrate the timescale of the prometaphase model by comparing the diffusion of a free bead to the diffusion of a similarly sized sphere in water. Using the Einstein-Stokes relation we calculate the diffusion coefficient of a bead $D = K_BT/6\pi\eta r$ as that of a sphere of radius $r = 0.048\mu m$ immersed in water at 298.15K. Using the viscosity of water $\eta = 8.9 \times 10^{-4} Pa \cdot s$ we obtain the diffusion coefficient of $5.12\mu m^2/s$.

We calculate the diffusion constant in reduced units using Einstein's relation. For a Langevin integrator with an inverse damping time of $\gamma = 0.1\frac{1}{\tau}$ and beads of mass $m = 10(\frac{\epsilon\tau^2}{\sigma^2})$, we get a mobility $\mu = v_d/(\gamma * m * v_d) = 1/(0.1\frac{1}{\tau} * 10(\frac{\epsilon\tau^2}{\sigma^2}) = 1\frac{\sigma^2}{\tau\epsilon}$ and a diffusion constant of $D = \mu K_BT = 1\frac{\sigma^2}{\tau}$ Comparing these two estimates of the diffusion constant, we obtain that the unit of time $\tau$ in our prometaphase simulations corresponds to approximately 1.8 milliseconds.

As an estimate, we will assume that the interphase DT40 chromosome's length scale is comparable to that measured for GM06990. Then, by the same calculation with a radius of $r = .0824\mu m$, we obtain that the unit of time $\tau$ in our interphase simulations corresponds to approximately 2.7 milliseconds.

### VII. CONSTANT-DISTANCE MEASUREMENT JUSTIFICATION

In "Constant-Distance" (CD) simulations, we restrict the chromosome's reaction coordinate to a small window around a reference distance $x_0$ using a harmonic potential $\boldsymbol{U}_{CD} = \frac{1}{2}k_p(\xi - \xi_0)^2$ given in the main text. We also claim in the main text that force at $\xi_0$ can be measured by taking an ensemble average of the force exerted by this pulling potential:

$$\mathbf{F}_{CF}(\xi_0) = \langle -k_p(\xi - \xi_0) \rangle \tag{33}$$

Here, we justify this claim:

We will consider the chromosome's macrostate to be defined by its reaction coordinate (extension) $\xi$. In the absence of the pulling potential, the Helmholtz free-energy of the chromosome is given by:

$$A(\xi) = U(\xi) - TS(\xi) \tag{34}$$

Taylor-expanding the Helmholtz to first order about $\xi_0$, we obtain:

$$A(\xi) \approx A(\xi_0) + \frac{\partial A(\xi)}{\partial \xi}|_{\xi=\xi_0} * (\xi - \xi_0) = A(\xi_0) + F(\xi_0) * (\xi - \xi_0) \tag{35}$$

Here, the sign of $F(\xi_0)$ is chosen so that positive forces correspond to stretching of the chromosome, and negative forces correspond to compression.

Applying $\boldsymbol{U}_{CD}$ effectively modifies the Helmholtz free energy as follows:

$$\begin{aligned} A_{CD}(\xi) &\approx A(\xi_0) + F(\xi_0) * (\xi - \xi_0) + \frac{1}{2}k_p(\xi - \xi_0)^2 \\ &= A(\xi_0) - \frac{F(\xi_0)^2}{2k_p} + \frac{1}{2}k_p\left(\xi - \xi_0 + \frac{F(\xi_0)}{k_p}\right)^2 \end{aligned} \tag{36}$$

where in the last step we have completed the square.

In equilibrium, the reaction coordinate $\xi$ will be distributed according to a Boltzmann distribution:

$$p(\xi) \propto \exp\left[-\beta A(\xi)\right] \propto \exp\left[-\beta \frac{1}{2} k_p \left(\xi - \xi_0 + \frac{F(\xi_0)}{k_p}\right)^2\right] \quad (37)$$

Thus, we find that $\xi$ is Gaussian distributed in equilibrium with mean $\langle \xi \rangle = \xi_0 - \frac{F(\xi_0)}{k_p}$ and (small) variance $\langle (\xi - \langle \xi \rangle)^2 \rangle = (\beta k_p)^{-1}$ The validity of this first-order approximation can be verified retroactively by checking that the distribution of $\xi$ follows this Gaussian distribution. Fitting a histogram of $\xi$ to a Gaussian distribution, we find excellent agreement with expected values of mean and variance (see figure S6C).

### VIII. MEASURING EXTENSION IN CONSTANT-FORCE ENSEMBLES

In this study, we employ two methods of force measurement: constant-force (CF) and constant-distance (CD) measurements. In the former, we generate an ensemble of structures with fixed end-to-end distance $\xi$. The thermodynamic potential which governs the behavior of such an ensemble is the Helmholtz Free Energy (see equation 34 above). The measured force is then given by $\frac{dA(\xi)}{d\xi}$. In the latter, we generate an ensemble of structures in the presence of a constant force $F$ which acts on the end-to-end distance, defined by the pulling potential $U_{CD} = -F\xi$. In this case, force is chosen and a histogram of extensions is calculated from the resulting ensemble of structures (see figure S3A). However, to obtain a force-extension curve we must map this extension histogram to a single extension. Two possible ways of doing this are to set the extension to the ensemble's mean extension ($\xi(F) = \langle \xi \rangle_F$), or to set the extension to it's most likely value ($\xi(F) = \text{argmax}(P(\xi)_F)$). Here $\xi = \langle \cdot \rangle_F$ represents a mean over the ensemble at constant force $F$ and $P(\xi)_F$ represents the probability density function of the reaction coordinate $\xi$ under constant force $F$.

Note that in the thermodynamic limit of infinite system size, the probability distribution $P(\xi)_F$ will converge to a Gaussian distribution, for which these two methods are equivalent. However, when dealing with systems of finite size there is no such guarantee, and the histograms in figure S3A are visibly asymmetric.

Here, we will show that in order to obtain a force-extension curve that matches the CD force-extension curve, we must select the maximum-likelihood extension. We start by rewriting the Gibbs free energy in terms of the Helmholtz free energy:

$$G(\xi) = A(\xi) - F\xi \quad (38)$$

Differentiating with respect to the extension $\xi$, we get:

$$\frac{dG(\xi)}{d\xi} = \frac{dA(\xi)}{d\xi} - F = F_{CD}(\xi) - F \quad (39)$$

We see that if $F(\xi) = F_{CD}$, then $\frac{dG(\xi)}{d\xi} = 0$. This implies that in order for the CF and CD force-extension curves to match, we must associate with the force F the value of $\xi$ which minimizes the Gibbs free energy. In the constant-force ensemble we also have that (up to a constant):

$$G(\xi) = -k_b T \log(P(\xi)) \quad (40)$$

Because the maximum-likelihood extension minimizes the Gibbs free energy, it is the correct choice of extension to pair with the applied force F. In prometaphse, the mean extensions and maximum-likelihood extensions are nearly identical. However, in interphase there is considerable divergence between the mean and maximum-likelihood extensions (see Fig. S3B). As is visible in figure 1, the maximum-likelihood extension matches well with the CD force-extension curve.

### IX. ORIENTATION CONSTRAINTS SIGNIFICANTLY ALTER INTERPHASE BUT NOT MITOTIC FORCE-EXTENSION BEHAVIOR

Conventionally, polymer conformation is described by the set of all coordinates of all beads $\vec{r} \in \mathbb{R}^{3N}$. To understand the force-extension behavior, we change coordinates to a so that the polymer conformation is described by the set $\{\mathbf{R}_{left}, \boldsymbol{\xi}, \vec{q}\}$, where $\vec{q} \in \mathbb{R}^{3N-6}$ describes the remaining degrees of freedom once the $\mathbf{R}_{left}$ and $\boldsymbol{\xi} = \mathbf{R}_{right} - \mathbf{R}_{left}$ have been specified. Because the potential energy function $U_{\text{MiChroM}}(\vec{r})$ is invariant to translations and rotations of the whole chromosome, we can rewrite its dependencies as $U_{\text{MiChroM}}(\vec{q}, \xi)$ where $\xi = |\boldsymbol{\xi}|$

$$Z_{\text{free}} = \int d^3\mathbf{R}_{left} d^3\boldsymbol{\xi} d^{3N-6}\vec{q} e^{-\beta U_{\text{MiChroM}}(\vec{q},\xi)} \quad (41)$$

$$= \int_0^\infty d\xi \left[V \times 4\pi\xi^2 \int d^{3N-6}\vec{q} e^{-\beta U_{\text{MiChroM}}(\vec{q},\xi)}\right] \quad (42)$$

$$= \int_0^\infty d\xi \left[Z_{\text{free}}(\xi)\right] \quad (43)$$

Where we have defined an extension-dependent partition function $Z_{\text{free}}(\xi) = V \times 4\pi\xi^2 \int d^{3N-6}\vec{q} e^{-\beta U_{\text{MiChroM}}(\vec{q},\xi)}$ accounting for the manifold of states with constant $\xi$.

Now we evaluate the partition function for a chromosome subjected to $U_{pin}(\mathbf{R}_{left}) + U_{slide}(\mathbf{R}_{right})$. To a good approximation, the effect of these potentials is to constrain $\boldsymbol{\xi}$ to point along the x axis.

$$Z_{\text{constrained}} = \int d^3\mathbf{R}_{left} d^3\boldsymbol{\xi} d^{3N-6}\vec{q}\delta^3(\mathbf{R}_{left})\delta(\boldsymbol{\xi}\cdot\boldsymbol{e_y})\delta(\boldsymbol{\xi}\cdot\boldsymbol{e_z})e^{-\beta U_{\text{MiChroM}}(\vec{q},\xi)} \tag{44}$$

$$= \int_0^\infty d\xi \left[\int d^{3N-6}\vec{q}e^{-\beta U_{\text{MiChroM}}(\vec{q},\xi)}\right] \tag{45}$$

$$= \int_0^\infty d\xi \left[Z_{\text{constrained}}(\xi)\right] \tag{46}$$

Where we have defined

$$Z_{\text{constrained}}(\xi) = \int d^{3N-6}\vec{q}e^{-\beta U_{\text{MiChroM}}(\vec{q},\xi)}$$

.

We now relate the partition functions for the free and constrained cases:

$$Z_{\text{Free}}(\xi) = V \times 4\pi\xi^2 Z_{\text{Constrained}}(\xi)$$

The extension-dependent Helmholtz free energies are given by:

$$A_{\text{constrained}}(\xi) = -\frac{1}{\beta}\log Z_{\text{constrained}}(\xi)$$

$$A_{\text{Free}}(\xi) = -\frac{1}{\beta}\log Z_{\text{Free}}(\xi) \tag{47}$$

$$= -\frac{1}{\beta}\log(4\pi V) - \frac{2}{\beta}\log(\xi) - \frac{1}{\beta}\log Z_{\text{Constrained}}(\xi) \tag{48}$$

$$= -\frac{1}{\beta}\log(4\pi V) - \frac{2}{\beta}\log(\xi) + A_{\text{Constrained}}(\xi) \tag{49}$$

Differentiating both sides with respect to $\xi$ and using $F(\xi) = \frac{\partial}{\partial\xi}A(\xi)$, we arrive at:

$$F_{\text{Free}}(\xi) = F_{\text{Constrained}}(\xi) - \frac{2}{\beta\xi} \tag{50}$$

This tells us that orientation constraints increase the force-extension curve by a "rotational" force contribution $F_{\text{Rot}}(\xi) = -\frac{2}{\beta\xi}$. This rotational force contributes a slope $k_{\text{rot}} = \frac{d}{d\xi}F_{\text{Rot}}(\xi)|_{\xi=\xi_0} = \frac{2}{\beta\xi_0^2}$ to the spring constant. We compare the magnitude of this contribution to the chromosome spring constants $k_{\text{sim}}$ at the native length measured in simulations:

| | Interphase | Mitosis |
|---|---|---|
| $\xi_0$ | $5.9\sigma$ | $13\sigma$ |
| $k_{\text{rot}}$ | $.06\epsilon/\sigma^2$ | $.01\epsilon/\sigma^2$ |
| $k_{\text{sim}}$ | $.09\epsilon/\sigma^2$ | $.73\ \epsilon/\sigma^2$ |

We see that in interphase, $k_{\text{rot}} \approx k_{\text{sim}}$, meaning that orientation constraints significantly alter the interphase chromosome's force-extension curve (and decreases its native length). However, in mitosis $k_{\text{rot}} << k_{\text{sim}}$, meaning that orientation constraints have little impact on the force-extension curve.

A

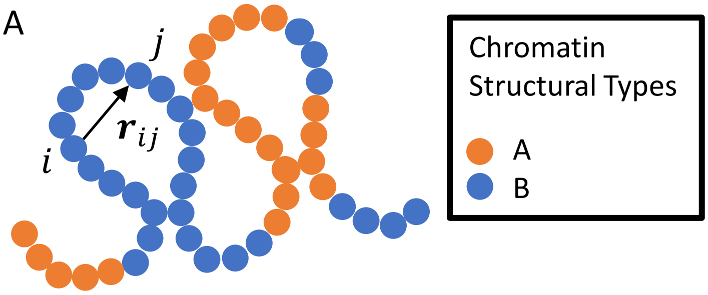


B

| Phase | $\alpha_{AA}$ | $\alpha_{BB}$ | $\alpha_{AB}$ |
|---|---|---|---|
| Interphase | -0.131 | -0.153 | -0.079 |
| Mitosis | -0.016 | -0.015 | -0.002 |

MiChroM Potential Function

Homopolymer:
$$U_{\text{MiChroM}}(\vec{r}) = U_{HP}(\vec{r})$$

Type-Type:
$$+ \sum_{\substack{k \geq l \\ k,l \in \text{ Types}}} \alpha_{kl} \sum_{\substack{i \in \{\text{ Loci of Type } k\} \\ j \in \{\text{ Loci of Type } l\}}} f(r_{ij})$$

Ideal Chromosome:
$$+ \sum_{d=3}^{500} \gamma(d) \sum_{i} f(r_{i,i+d})$$

$$f(r_{ij}) = \frac{1}{2}(1 + \tanh[\mu(r_c - r_{ij})])$$

C

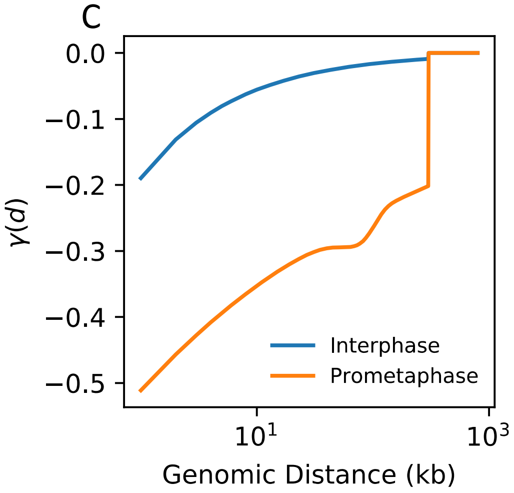


Figure S1: Model Details

a) Chromosomes are modeled as a string of beads, each bead representing 50 kb. Beads are assigned an epigenetic type A or B, corresponding roughly to euchromatin and heterochromatin. Beads interact through a homopolymer potential, as well as through pairwise “type-type” interactions that depend on the structural types of the interacting beads and an “ideal chromosome” interaction that depends on the genomic distance between interacting beads. During training, the type-type and ideal chromosome interaction parameters are tuned to obtain the best agreement between simulated and experimental Hi-C contact maps.

b) Type-Type interaction parameters for the interphase and prometaphase chromosome. When two beads come within a cutoff distance of each other, they experience a free energy loss that depends on the structural types of the interacting beads.

c) The Ideal Chromosome interaction parameters for interphase and prometaphase chromosomes. When two beads come within a certain cutoff distance of each other, they experience a free energy loss that depends on the genomic distance (contour length) that separates them. The depth of this free energy loss as a function of genomic distance, $\gamma(d)$, is shown here for both interphase (blue) and mitosis (orange).

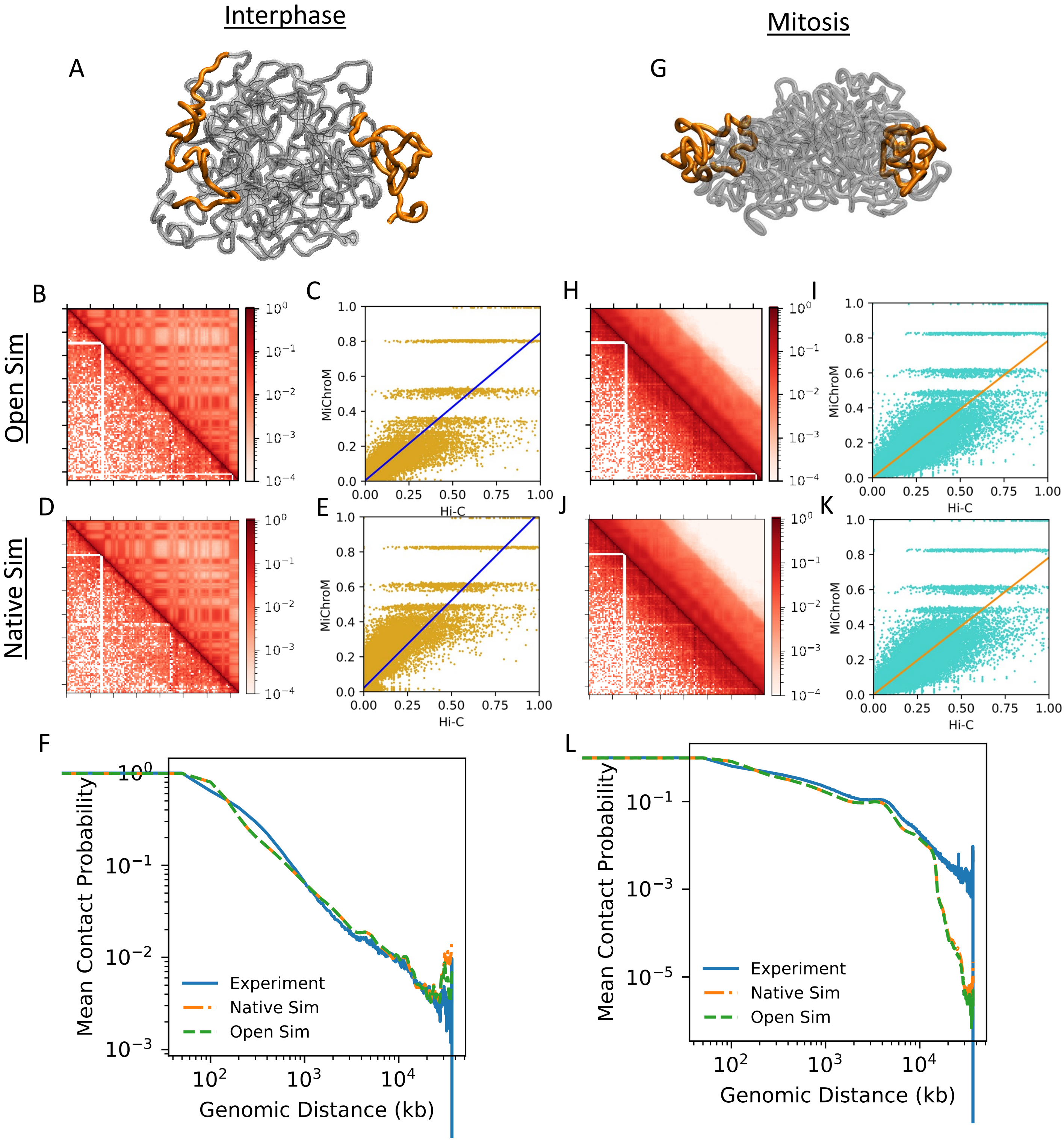

Interphase
Mitosis
A
G
B
C
H
I
Open Sim
D
E
J
K
Native Sim
MiChroM
Hi-C
F
L
Mean Contact Probability
Genomic Distance (kb)
Experiment
Native Sim
Open Sim

Figure S2: Comparison between model and experiment.
(A/G) Snapshot of a simulated interphase/mitotic chromosome structure. Pull groups consisting of the first and last 50 beads (2500 kb) are highlighted in orange.
(B/H) Comparison between simulated (upper right) and experimental (lower left) contact maps. Contact maps were generated form "Open" simulations, in which the chromosomes are simulated with no pulling or orientational constraints.
(C/I) Scatter plot of experimental Hi-C contact probabilities and simulated contact probabilities from "Open" simulations. Pearson's correlations $r = .91$ and $r = .90$ were measured for interphase and mitosis respectively.
(D/J) Comparison between simulated (upper right) and experimental (lower left) contact maps. Contact maps were generated form "native" simulations, in which the chromosome's left pull group is constrained to the origin and it's right pull group is constrained to the x-axis but allowed to slide along it.
(E/K) Scatter plot of experimental Hi-C contact probabilities and simulated contact probabilities from "Native" simulations. Pearson's correlations $r = .87$ and $r = .90$ were measured for interphase and mitosis respectively.
(F/J) Comparison between simulated and experimental contact scaling curves. Blue line shows experimental contact scaling curve. Dotted green line shows simulated contact scaling curve for "open" simulations. Dash-dotted orange line shows contact scaling curve for "native" simulation. Simulated and experimental maps are in good agreement. Notably, in interphase, there is an increase in contacts between pull groups for the "native" simulation relative to the "open" simulation. The apparent disagreement between simulated and experimental contacts at long genomic distance in prometaphase is an artifact of the log scale, as the contact frequency for both is very small.

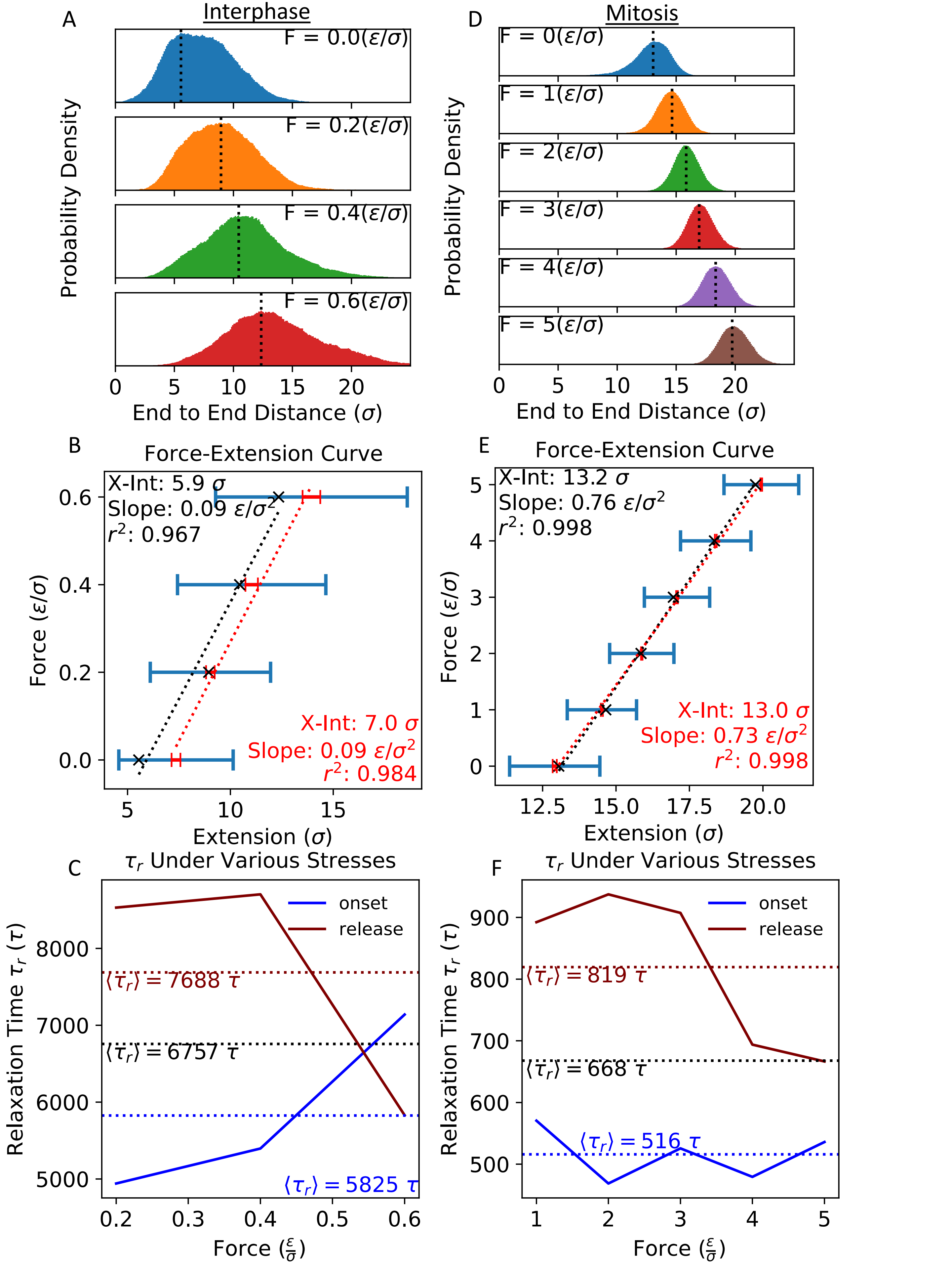

A
Interphase
F = 0.0(ε/σ)
F = 0.2(ε/σ)
F = 0.4(ε/σ)
F = 0.6(ε/σ)
Probability Density
End to End Distance (σ)
D
Mitosis
F = 0(ε/σ)
F = 1(ε/σ)
F = 2(ε/σ)
F = 3(ε/σ)
F = 4(ε/σ)
F = 5(ε/σ)
Probability Density
End to End Distance (σ)
B
Force-Extension Curve
X-Int: 5.9 σ
Slope: 0.09 ε/σ²
r²: 0.967
X-Int: 7.0 σ
Slope: 0.09 ε/σ²
r²: 0.984
Force (ε/σ)
Extension (σ)
E
Force-Extension Curve
X-Int: 13.2 σ
Slope: 0.76 ε/σ²
r²: 0.998
X-Int: 13.0 σ
Slope: 0.73 ε/σ²
r²: 0.998
Force (ε/σ)
Extension (σ)
C
τ_r Under Various Stresses
onset
release
⟨τ_r⟩ = 7688 τ
⟨τ_r⟩ = 6757 τ
⟨τ_r⟩ = 5825 τ
Relaxation Time τ_r (τ)
Force (ε/σ)
F
τ_r Under Various Stresses
onset
release
⟨τ_r⟩ = 819 τ
⟨τ_r⟩ = 668 τ
⟨τ_r⟩ = 516 τ
Relaxation Time τ_r (τ)
Force (ε/σ)

Figure S3: Chromosome Behavior Under Constant Force.
(A/D) Histograms of the interphase/mitotic chromosome extension distance during constant-force sampling at each force sampled. Vertical dotted lines mark the extensions of maximum likelihood.
(B/D) Force-Extension curve calculated from constant-force sampling. Blue error bars show standard deviation of the extension for each force. Red error bars show the standard error of the mean extension when averaged over 40 replica trajectories. Black x-marks show maximum-likelihood extension. Linear fits for both the mean extensions and maximum-likelihood extensions are shown as black and red dotted lines, respectively.
(C/E) The relaxation times determined by exponential fitting of the mean extension time-courses in main text fig. 2. Blue shows the relaxation time measured after the sudden onset of a constant force. The maroon line shows the relaxation time measured after the sudden release of the force. Horizontal dotted lines show the mean relaxation times over force onset measurements, force release measurements, and all measurements (black).

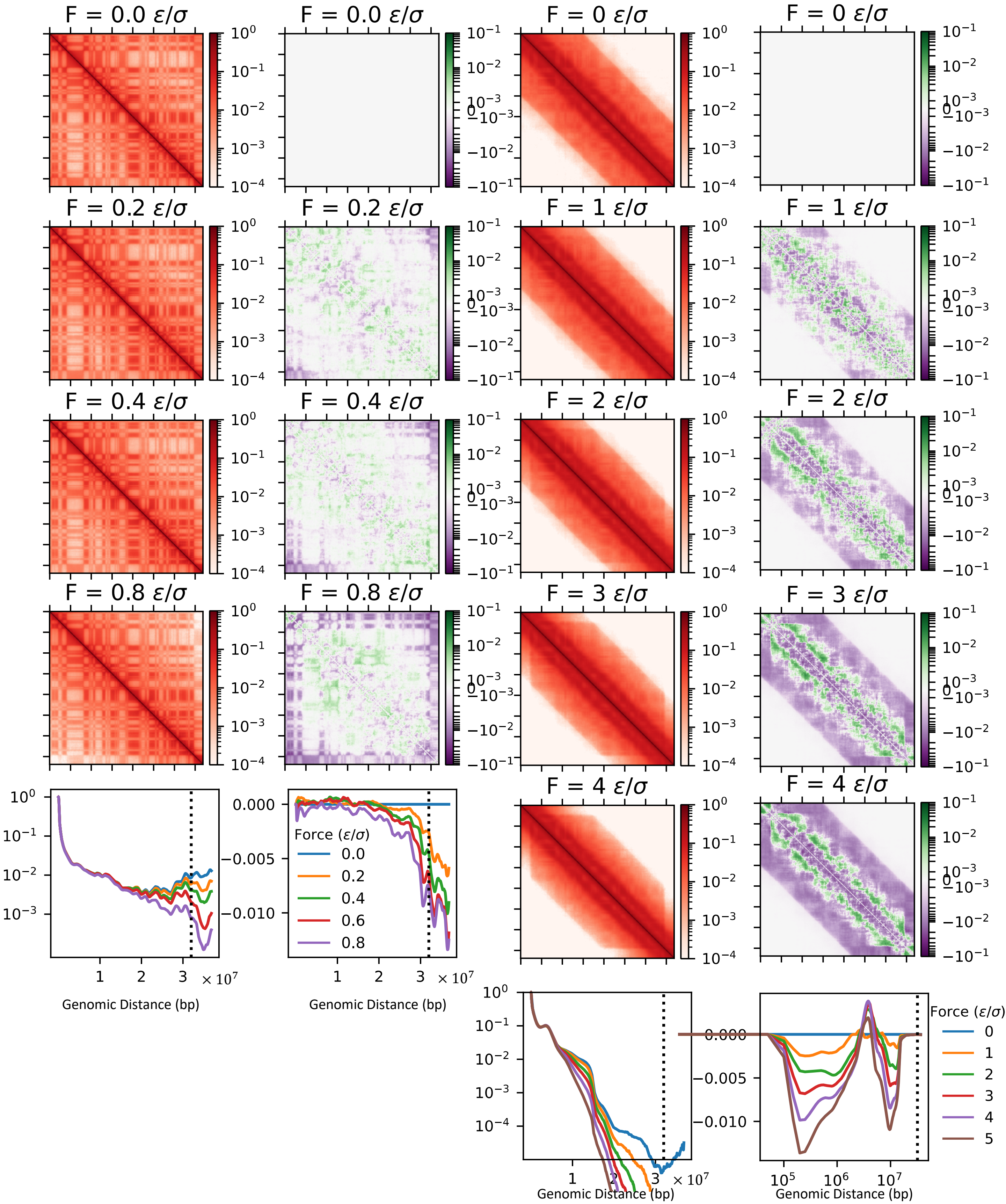
F = 0.0 ε/σ
F = 0 ε/σ
F = 0.2 ε/σ
F = 1 ε/σ
F = 0.4 ε/σ
F = 2 ε/σ
F = 0.8 ε/σ
F = 3 ε/σ
F = 4 ε/σ
Force (ε/σ)
0.0
0.2
0.4
0.6
0.8
Genomic Distance (bp)
× 10^7
Force (ε/σ)
0
1
2
3
4
5

Figure S4: Simulated contact maps of interphase (left) and mitosis (right) with constant applied force. Red contact maps show absolute contact probability, plotted in log scale. Green and purple contact maps show contact probability relative to zero-force ensemble, plotted with a symmetric log scale with linear cutoff $10^{-3}$. Here, green indicates an increase in contact probability under tension and purple indicates a decrease in contact probability under tension. Below each column, the maps are averaged over their diagonals to give the absolute and relative contact scaling curves. In scaling curves, vertical the dotted line at 639 beads marks the genomic distance beyond which all contacts are between the two pull groups. In interphase, pulling mainly disrupts contacts between the pull groups. In prometaphase, pulling has almost no effect on the pull groups' interactions, but leads to a distinct pattern of lost contacts around 200 kb and 10000 kb as well as a slight enhancement of interactions around 4000 kb

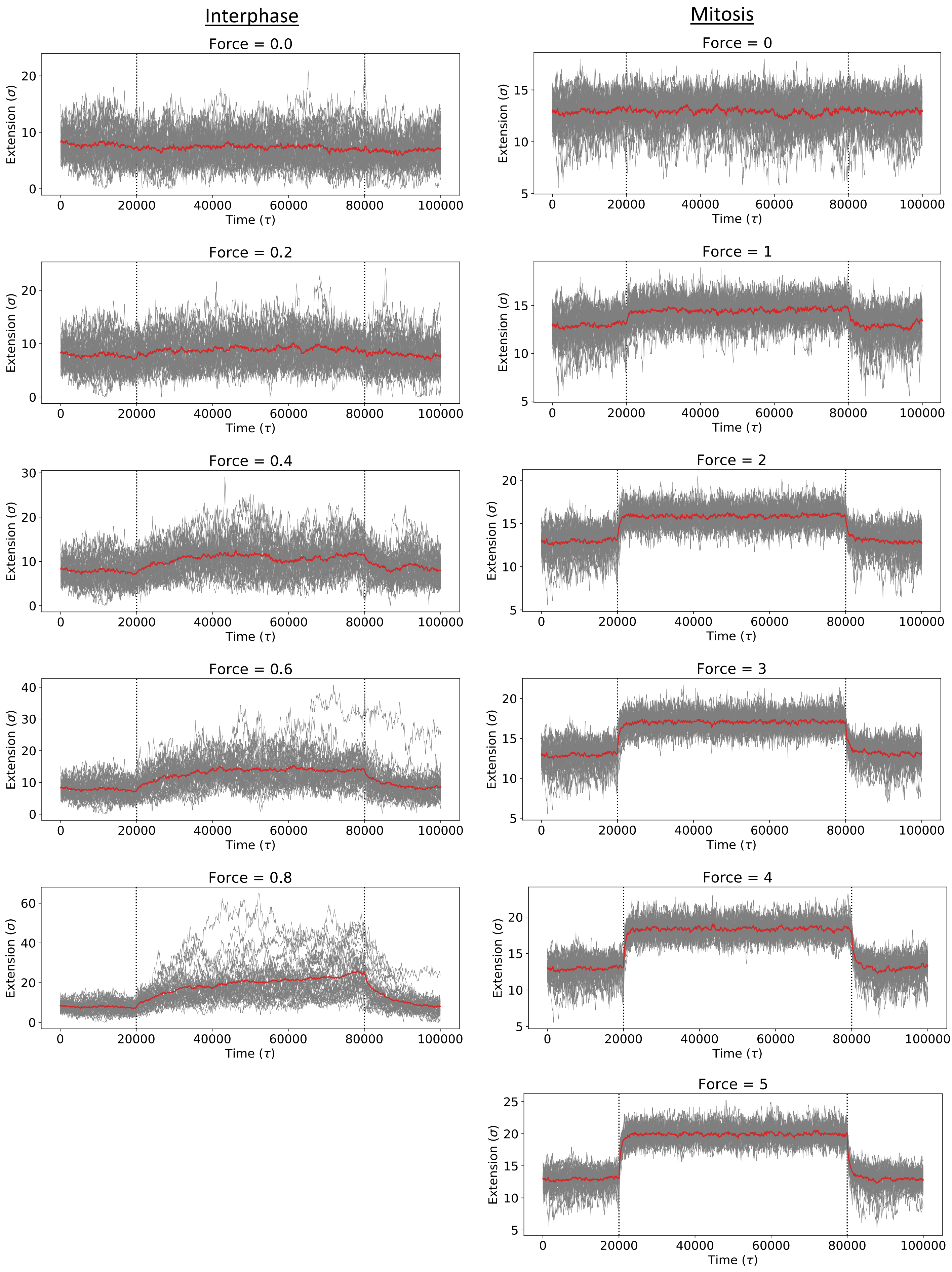

Interphase
Mitosis
Force = 0.0
Force = 0
Force = 0.2
Force = 1
Force = 0.4
Force = 2
Force = 0.6
Force = 3
Force = 0.8
Force = 4
Force = 5
Extension (σ)
Time (τ)
0
20000
40000
60000
80000
100000

Figure S5: Individual Chromosome trajectories during simulated stress-relaxation experiment for interphase (left column) and mitosis (right column). In the stress-relaxation experiment, a constant force was suddenly applied to the chromosome's pull groups at time 20000 $\tau$, then released at time 80000 $\tau$. To reduce noise, the chromosome's extension was averaged over 40 trajectories. The individual trajectories are shown here in gray, and their average in red (as well as main text fig. 2) .

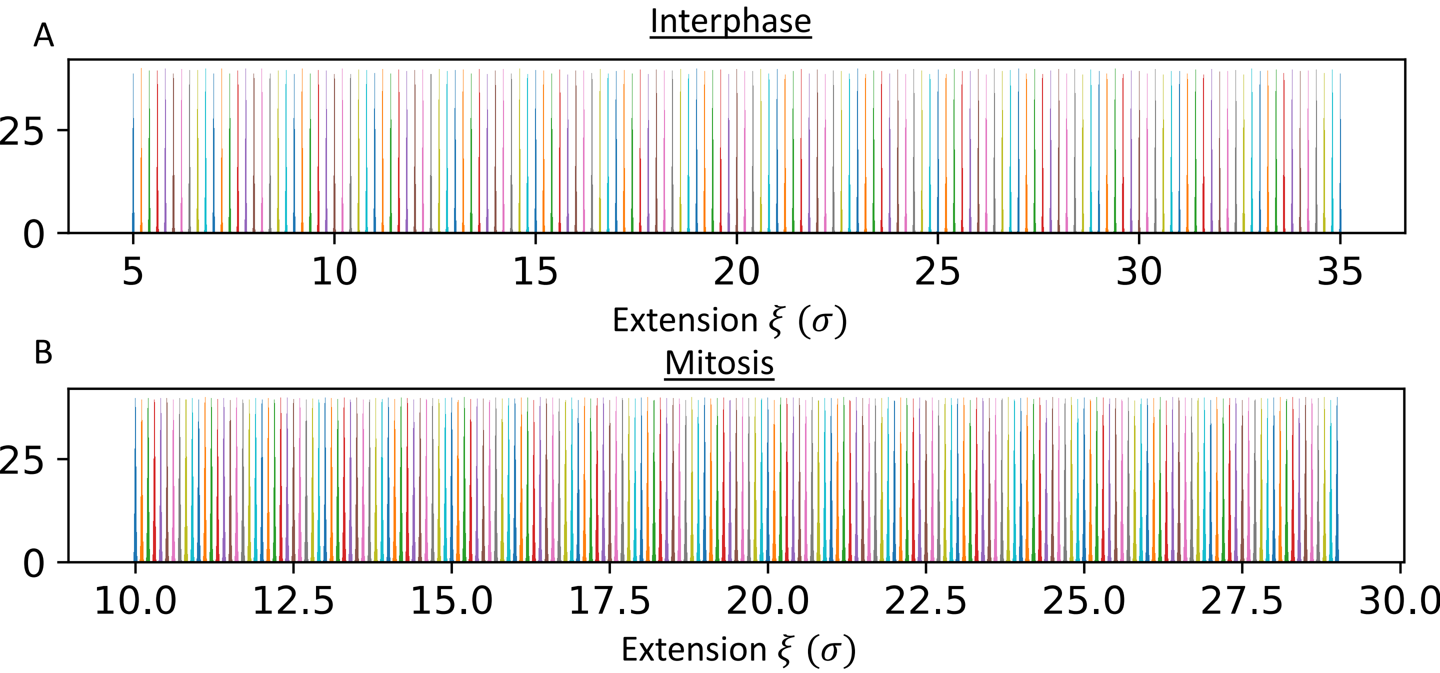


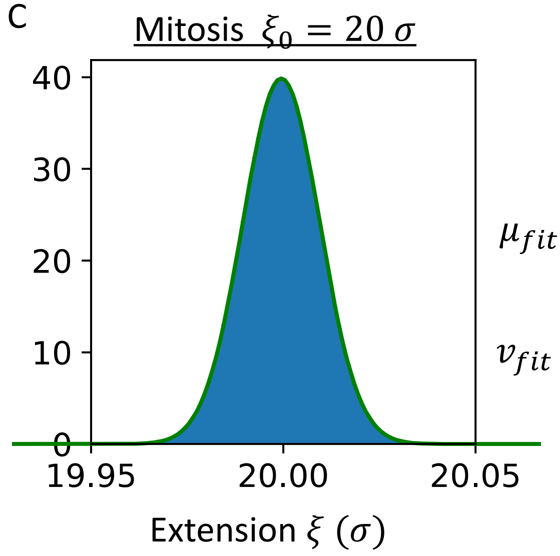


$$P(\xi) = \frac{1}{2\pi v^2} e^{\frac{(x-\mu)^2}{2v^2}}$$

$\mu_{fit} = 19.999486\ \sigma$ $\quad \mu_{expected=}\ \xi_0 - \frac{F(\xi_0)}{k_p} = 19.999487\ \sigma$

$v_{fit} = 0.010006\ \sigma$ $\quad v_{expected} = \left(\beta k_p\right)^{-\frac{1}{2}} = 0.01\ \sigma$

Fig S6: Histograms of CD extension windows in interphase and mitosis.
(A/B) Histograms of chromosome extension for each chosen reference distance during CD pulling of the interphase/mitotic chromosome. During CD pulling experiment, chromosome extension $\xi$ is restricted to a very small window about a reference distance $\xi_0$ with a strong harmonic potential $U_{CD} = \frac{1}{2} k_p (\xi - \xi_0)^2$ A large value $k_p = 10000\ \epsilon/\sigma^2$ is selected. Chosen reference distances are 5, 5.2, … 35 in interphase and 10, 10.1, …, 29 in mitosis.
(C) Detailed view of a single representative histogram of chromosome extension during CD pulling (mitosis, $\xi_0 = 20\ \sigma$). Green line shows fit to gaussian distribution, which is an excellent approximation to the distribution. To the right of the plot, fitted gaussian parameters are compared to those expected based on the first-order Taylor series expansion argument in the SI. Agreement between expected and fitted parameters indicates self-consistency of the perturbation used in the SI.

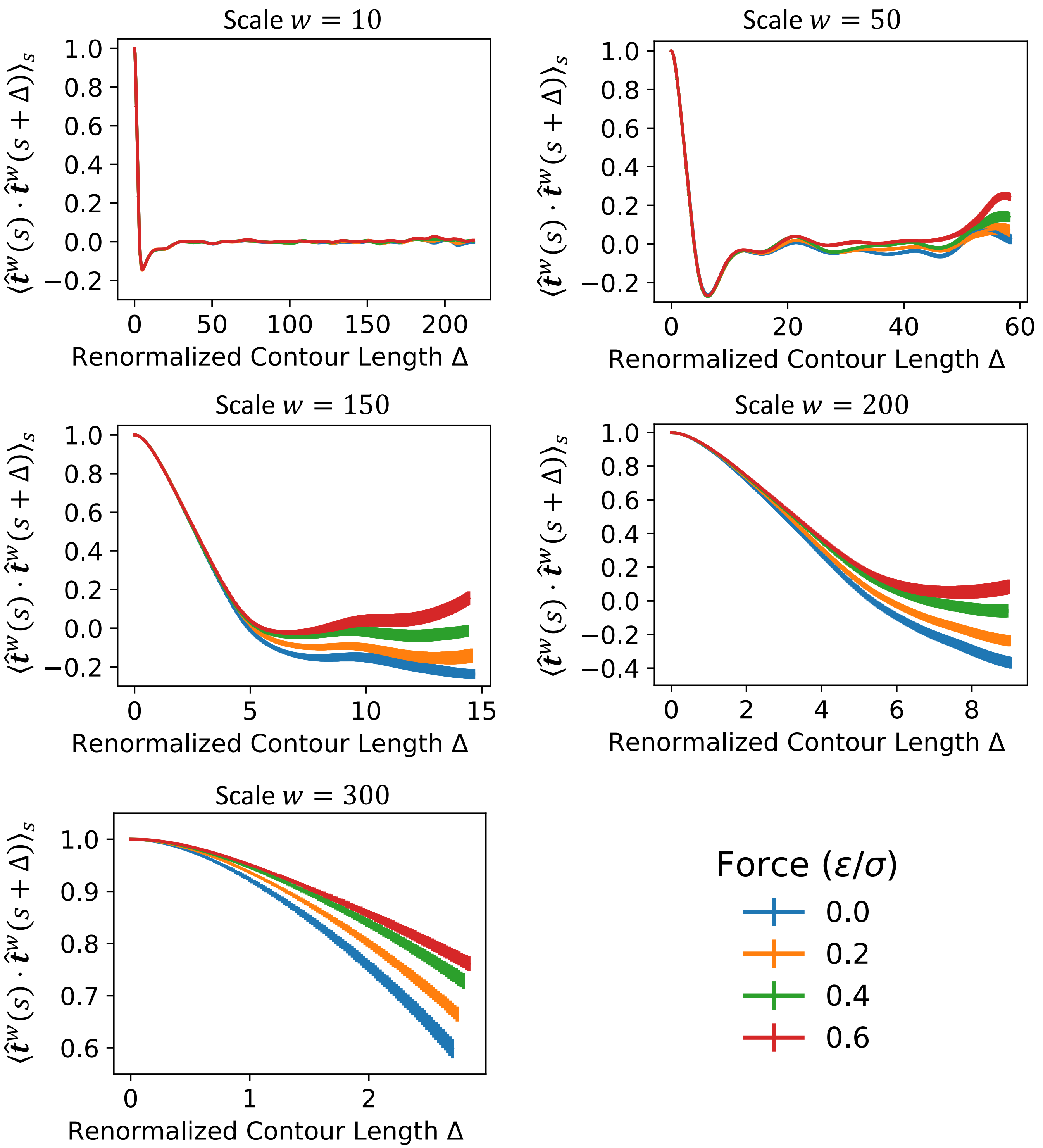


Figure S7: Renormalized tangent correlation functions $\tilde{T}^w(\Delta) = \langle \hat{\boldsymbol{t}}^w(s) \cdot \hat{\boldsymbol{t}}^w(s+\Delta) \rangle_s$ for interphase chromosome 7 under constant forces F = 0, 0.2, 0.4, 0.6. Vertical error bars show the standard error of the mean over 40 replica trajectories. Coarse-grained bead sizes 10, 50, 150, 200, and 300 are shown here.

Increasing the coarse-grained bead size effectively increases the length scale at which chromosome structure is analyzed. For small coarse-grained bead size of 10 beads x 50kb per bead, there is no visible change in the tangent correlation function, indicating that chromosome organization at this scale is unaffected. At larger coarse-grained bead sizes, the tangent correlation function increases with force, indicating the realignment of chromosome structure at larger scales.

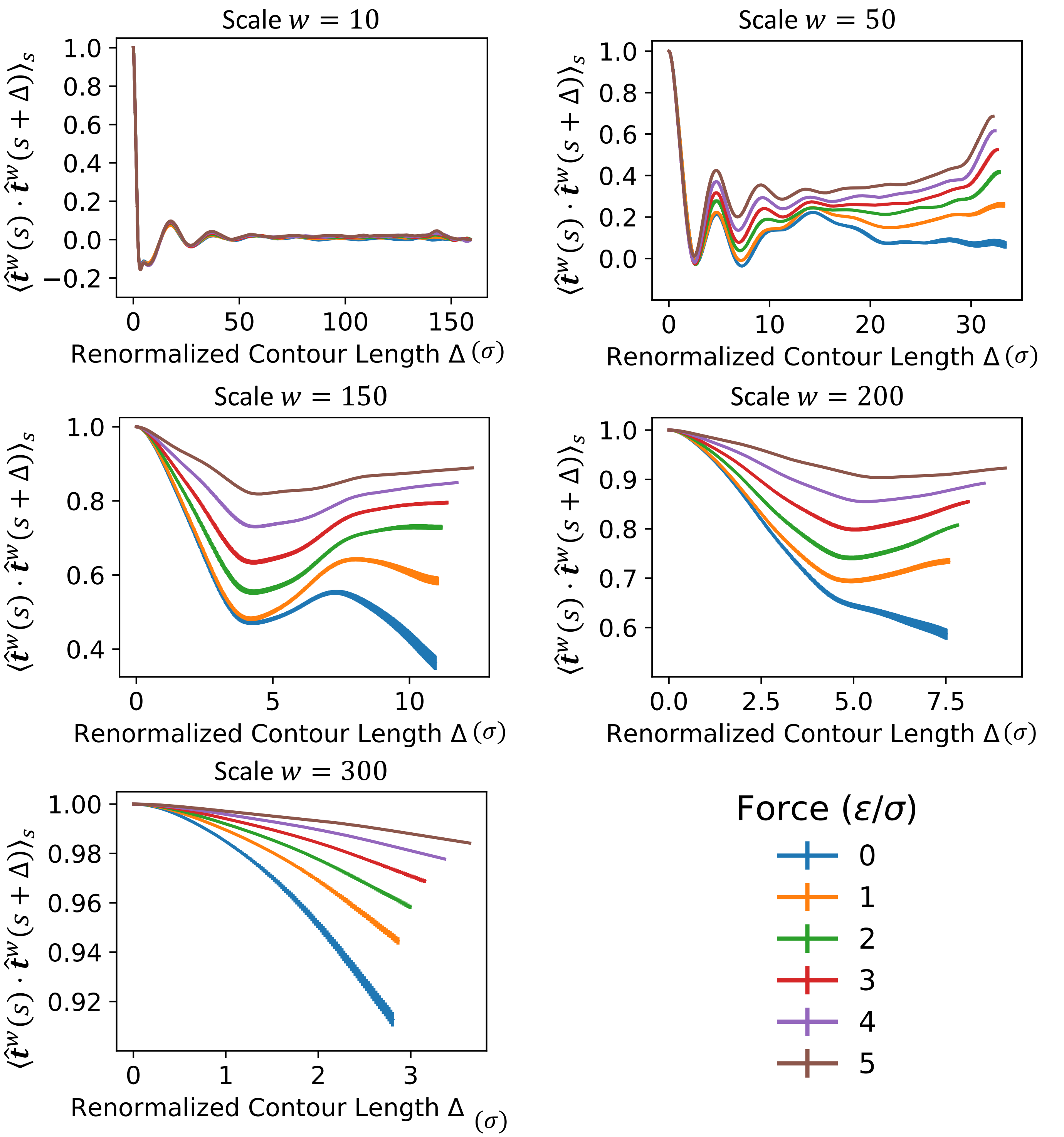


Figure S8: Renormalized tangent correlation functions $\tilde{T}^w(\Delta) = \langle \hat{\boldsymbol{t}}^w(s) \cdot \hat{\boldsymbol{t}}^w(s+\Delta) \rangle_s$ for interphase chromosome 7 under constant forces F = 0, 0.2, 0.4, 0.6. Vertical error bars show the standard error of the mean over 40 replica trajectories. Coarse-grained bead sizes 10, 50, 150, 200, and 300 are shown here. Vertical error bars show the standard error of the mean over 40 replica trajectories.

Coarse-grained bead sizes 10, 50, 150, 200, and 300 are shown here. Increasing the coarse-grained bead size effectively increases the length scale at which chromosome structure is analyzed. For small coarse-grained bead size of 10 beads x 50kb per bead, there is no visible change in the tangent correlation function, indicating that chromosome organization at this scale is unaffected. At larger coarse-grained bead sizes, the tangent correlation function increases with force, indicating the realignment of chromosome structure at larger scales.

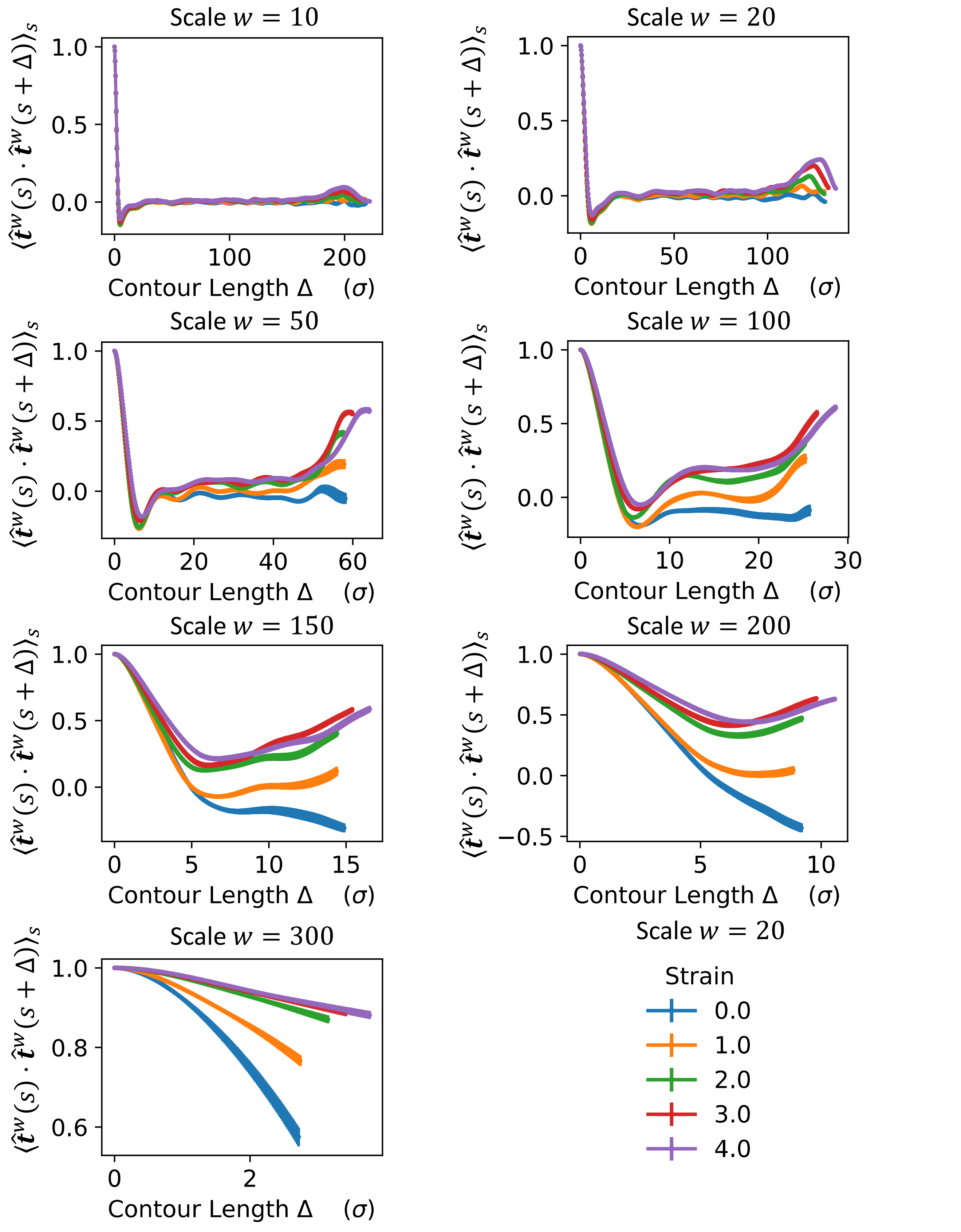


Figure S9: Renormalized tangent correlation functions $\tilde{T}^w(\Delta) = \langle \hat{\boldsymbol{t}}^w(s) \cdot \hat{\boldsymbol{t}}^w(s+\Delta) \rangle_s$ for interphase chromosome 7 for ensembles at fixed axial strains 0, 1, 2, 3, 4. Vertical error bars show the standard error of the mean over 40 replica trajectories. Coarse-grained bead sizes 10, 20, 50, 100, 150, 200, and 300 are shown here.

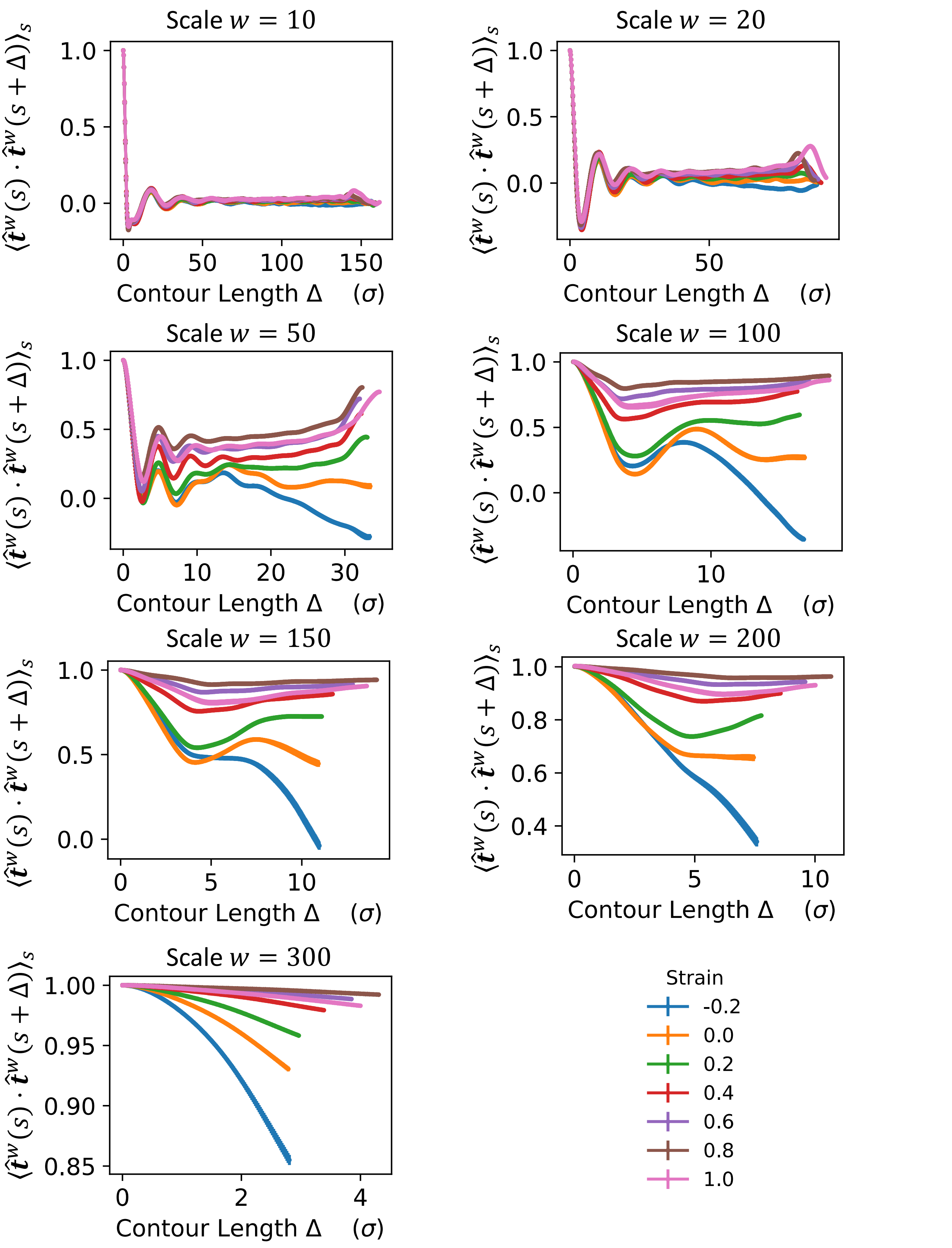


Figure S10: Renormalized tangent correlation functions $\tilde{T}^w(\Delta) = \langle \hat{\boldsymbol{t}}^w(s) \cdot \hat{\boldsymbol{t}}^w(s+\Delta) \rangle_s$ for mitotic chromosome 7 for ensembles at fixed axial strains -.2, 0, .2, .4, .6, .8, 1. Vertical error bars show the standard error of the mean over 40 replica trajectories. Coarse-grained bead sizes 10, 20, 50, 100, 150, 200, and 300 are shown here.

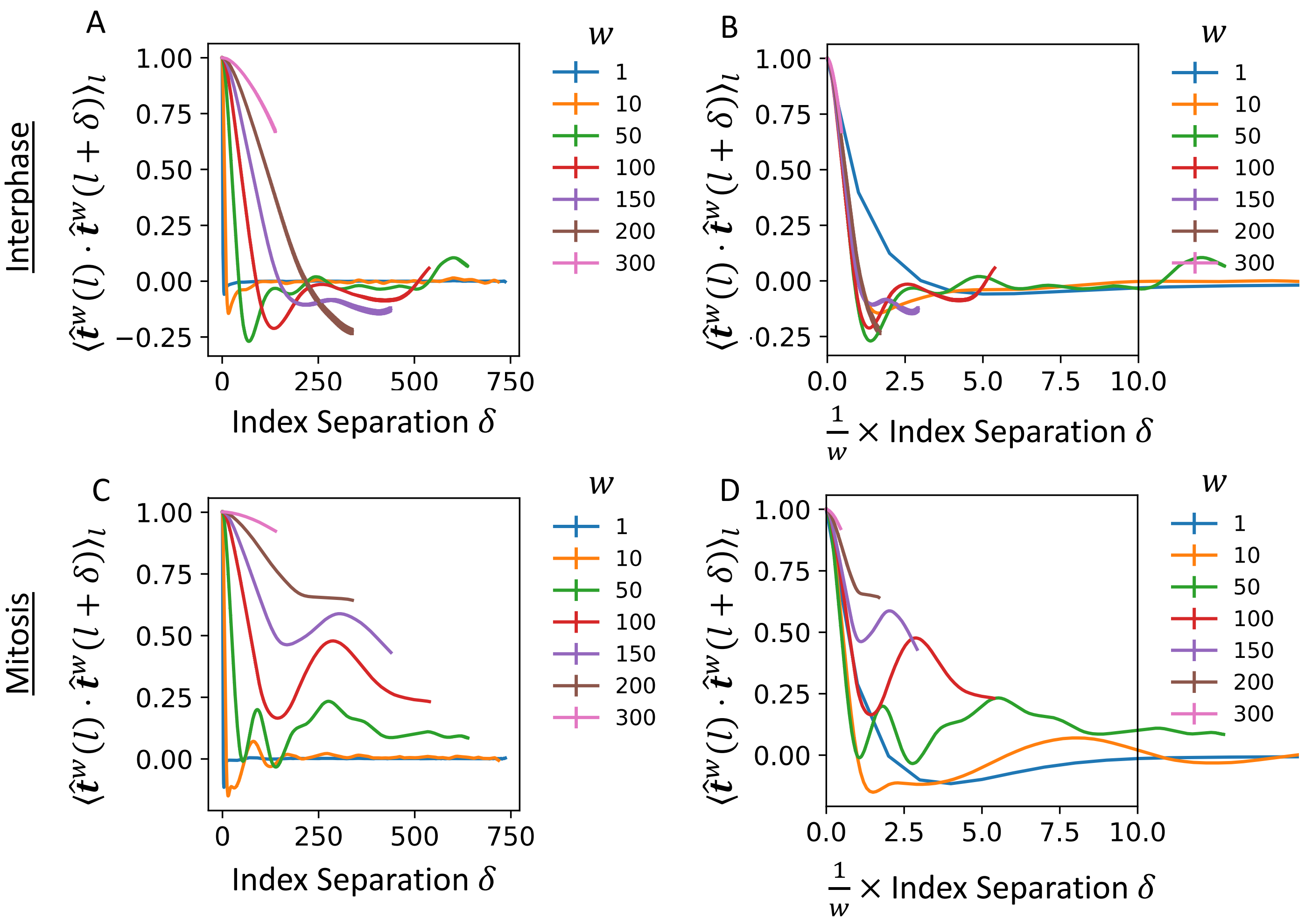


Figure S11: Structural scale invariance is destroyed in the transition from interphase to mitosis. (A/C) Renormalized tangent correlation functions $\tilde{T}^w(\delta) = \langle \hat{\boldsymbol{t}}_l{}^w \cdot \hat{\boldsymbol{t}}_{l+\delta}{}^w \rangle_l$ for interphase chromosome 7 in interphase/mitosis.

(B/D) Same plots as the left column, except that the x-axis is rescaled by the window size w. In interphase, this causes all correlation functions at at or above 10 beads to collapse into a single curve, with reasonable precision. In mitosis, rescaling of the x-axis does not cause such a collapse. This indicates a loss of scale-invariance upon the interphase-to-mitosis transition.

SI: Chromosome Radius, CG bead Rg and Linear Compaction vs. Extension

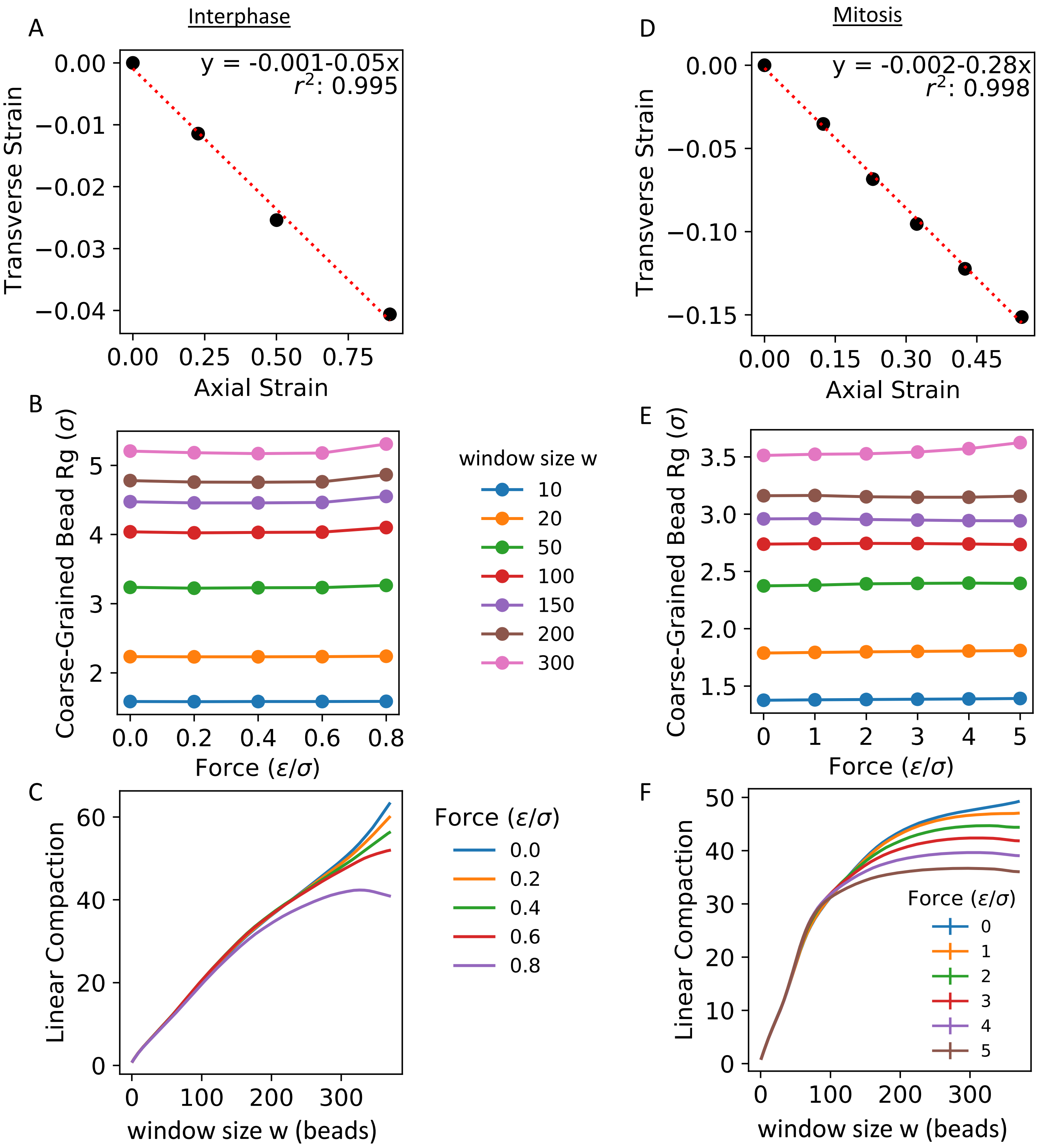

Figure S12: Other structural features of interphase (left) and prometaphase (right) chromosome under constant force:
(A/D) Normalized chromosome transverse strain vs. axial strain during CF pulling in interphase/mitosis. Linear fits give Poisson ratios of .05 and .28 for interphase and mitosis, respectively. These values can be compared to a Poisson ratio of .067 measured for mitotic chromosomes in Poirier et. al, *Molecular Biology of the Cell* (2000)
(B/E) Radius of gyration of collections of beads in sliding windows of various sizes as a function of force applied to chromosomes in CF pulling experiment. We observe that coarse-grained bead sizes are nearly constant as tension increases. This indicates that changes in chromosome structure are due to the rearrangement of the coarse-grained beads of the renormalized chain, rather than the deformation of the coarse-grained beads.
(C/F) Degree of linear compaction: $L/L'$ where L is the contour length of the bare polymer chain and L' is the contour length of the renormalized chain obtained by averaging bead positions over sliding windows of w beads. Linear compaction as a function of renormalization scale (sliding window size w) is shown for chromosomes under various forces. In interphase, linear compaction appears unaffected by applied tension. In prometaphase, linear compaction decreases under tension at large renormalization scales greater than 150 beads. This indicates that pulling causes chromosomes to stretch at large length scales without affecting structure at small scales.

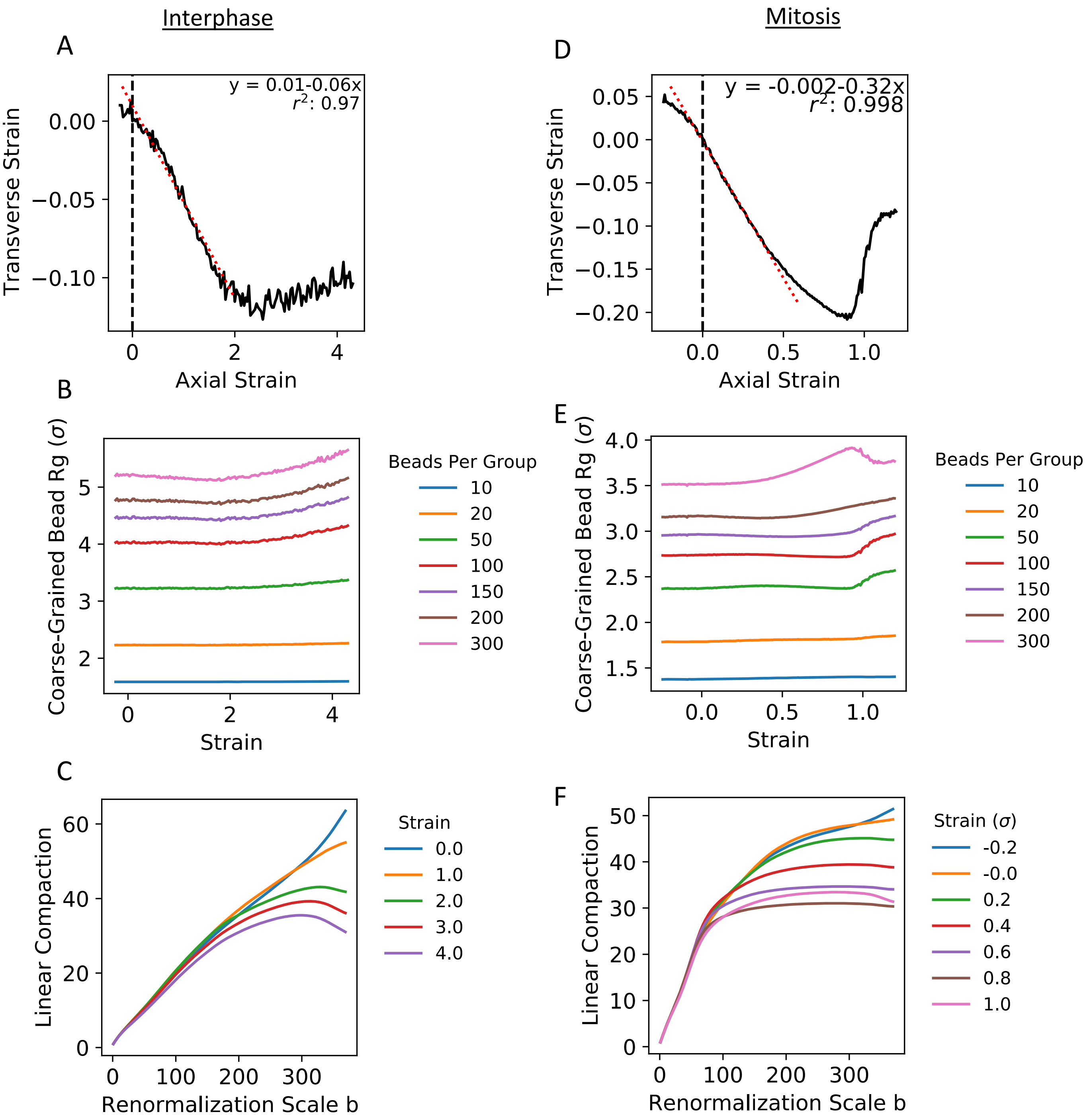


Fig S13: Radius-Extension, coarse-grained bead RG and Compaction vs. strain for CD simulations. (A/D) Interphase/Mitotic chromosome transverse strain vs. normalized mean chromosome extension during CF pulling in interphase/mitosis. Linear fits give Poisson ratios of .06 and .32 for interphase and mitosis, respectively. These values can be compared to a Poisson ratio of .067 measured for mitotic chromosomes in Poirier et. al, *Molecular Biology of the Cell* (2000).
(B/E) Radius of gyration of collections of beads in sliding windows of various sizes as a function of axial strain in CD pulling experiment.
(C/F) Degree of linear compaction as a function of renormalization scale (sliding window size w) as a function of strain in CD pulling experiment.

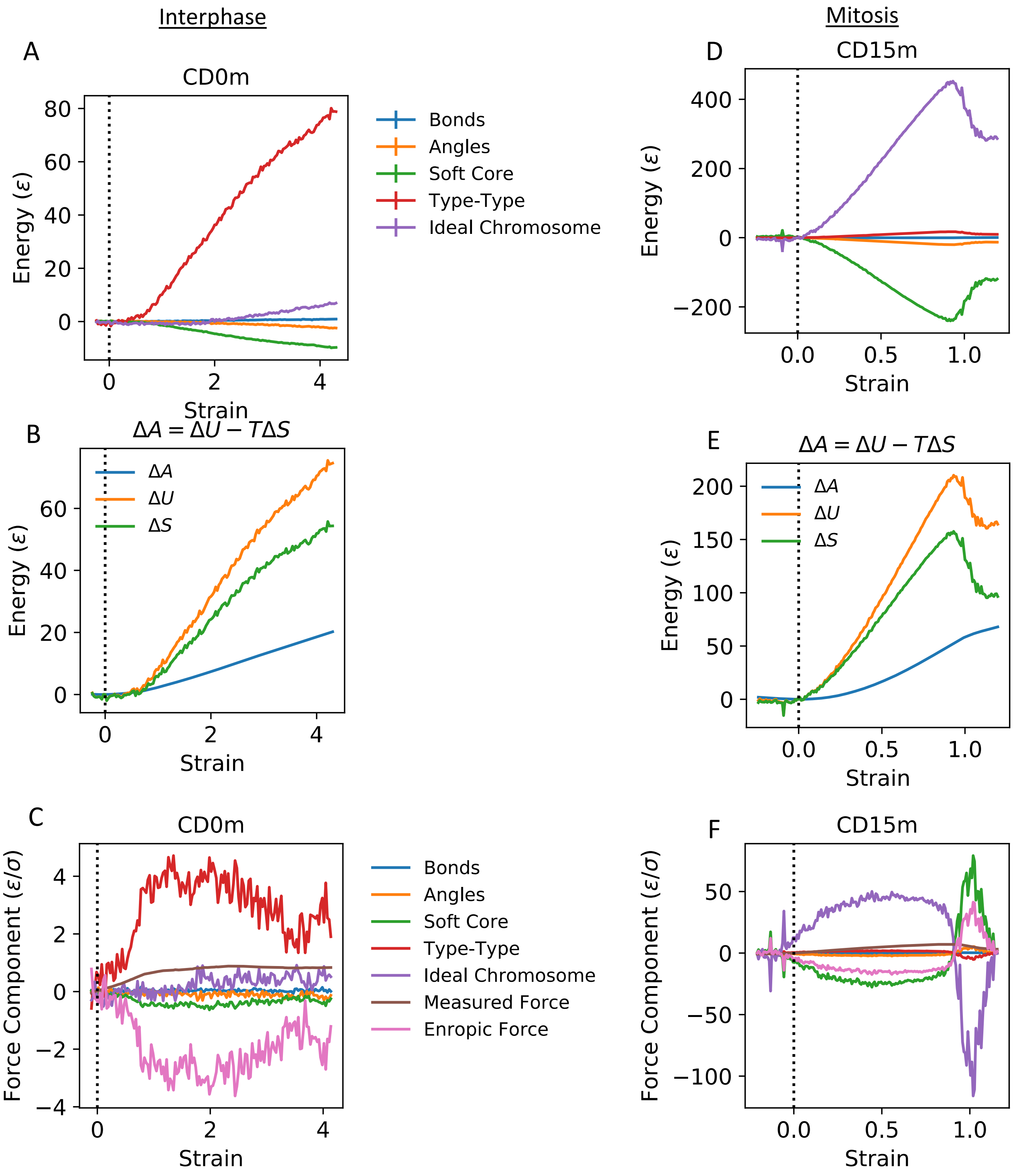

Interphase
Mitosis
A
CD0m
Energy (ε)
Strain
Bonds
Angles
Soft Core
Type-Type
Ideal Chromosome
D
CD15m
Energy (ε)
Strain
B
ΔA = ΔU − TΔS
ΔA
ΔU
ΔS
Energy (ε)
Strain
E
ΔA = ΔU − TΔS
ΔA
ΔU
ΔS
Energy (ε)
Strain
C
CD0m
Force Component (ε/σ)
Strain
Bonds
Angles
Soft Core
Type-Type
Ideal Chromosome
Measured Force
Enropic Force
F
CD15m
Force Component (ε/σ)
Strain

Figure S14: Potential Energy vs. Extension from constant-distance sampling for interphase (left) and prometaphase (right) chromosome:
(A/D) Interphase/Mitotic chromosome potential energy components vs. axial strain. In agreement with constant-force results, pulling primarily disrupts the interphase chromosome's type-type interactions are the prometaphase chromosome's ideal-chromosome interactions.
(B/E) Total potential energy, free energy, and entropy vs. mechanical strain. Total potential energy is calculated as the sum of the potential energy components in (A/D). Free energy is computed by numerical integration of the force-extension curve from constant-distance sampling. Entropy is calculated as the difference between total potential energy and free energy using the equation: $\Delta A = \Delta U - T\Delta S$. Here, T has been set to unity. Note that for the one-dimensional reaction coordinate, free energy and potential of mean force are equal. Also note that $\Delta E = \Delta \mathrm{U}$ as simulations are carried out at constant temperature so that mean kinetic energy is constant. We observe that potential energy increases faster than free energy as chromosomes stretch, indicating an increase in entropy as chromosomes unfold.
(C/F) Gradients of potential energy components, free energy, and entropy with respect to extension. This gives the "components" of the force associated with each potential energy component. Positive force components for the ideal chromosome and type-type interactions resist chromosome stretching. A negative force component for soft-core interaction favors unfolding. The negative force component attributed to entropy reflects the entropic favorability of stretching/unfolding.

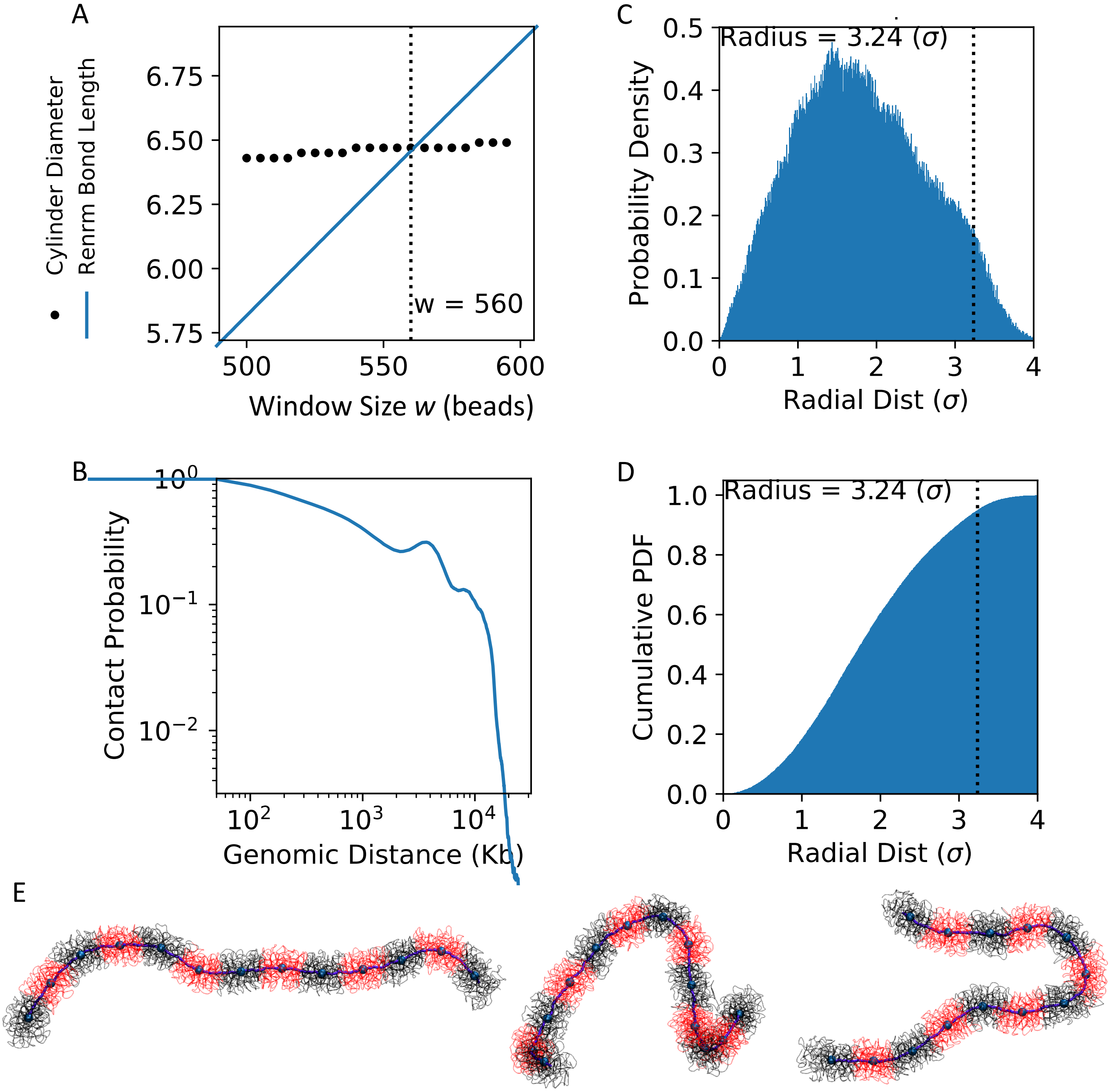


Figure S15: Calculation of Prometaphase Chromosome Persistence Length:

(A) Mitotic chromosome diameter $2R^w$, defined according to SI equation 12 and mean renormalized bond size $\langle|\,\tilde{\boldsymbol{v}}_i^w|\rangle_i$ defined according to SI equation 5 are shown as a function of renormalization scale w. A “natural” renormalization scale of w=560 is chosen so that these values agree.

(B) Contact Scaling Curve of “extended” mitotic chromosome 7 consisting of 10 copies of chromosome 7 concatenated head to tail.

(C) Histogram of the distances of chromosomal beads from the ”thin core” obtained by averaging over a sliding window of w=560 beads.

(D) Cumulative distribution calculated from histogram in (C). Vertical dotted line shows where cumulative distribution reaches 0.95, defined as the chromosome’s radius.

(E) Snapshots of extended mitotic chromosome structures. Renormalized Chain $\tilde{\boldsymbol{r}}_i^w = \frac{1}{w}\sum_{l=i}^{i+w} \boldsymbol{r}_i$ is shown in purple. One set of nearest non-overlapping neighbor renormalized beads $\{\tilde{\boldsymbol{r}}_1^w, \tilde{\boldsymbol{r}}_{1+w}^w, \tilde{\boldsymbol{r}}_{1+2w}^w, \ldots\}$ is shown as large blue spheres. Blue spheres are the centers of mass of the associated raw beads shown in alternating red and black thin tubing.

DT40 Chromosome 7 DT40 “Extended” Chromosome 7

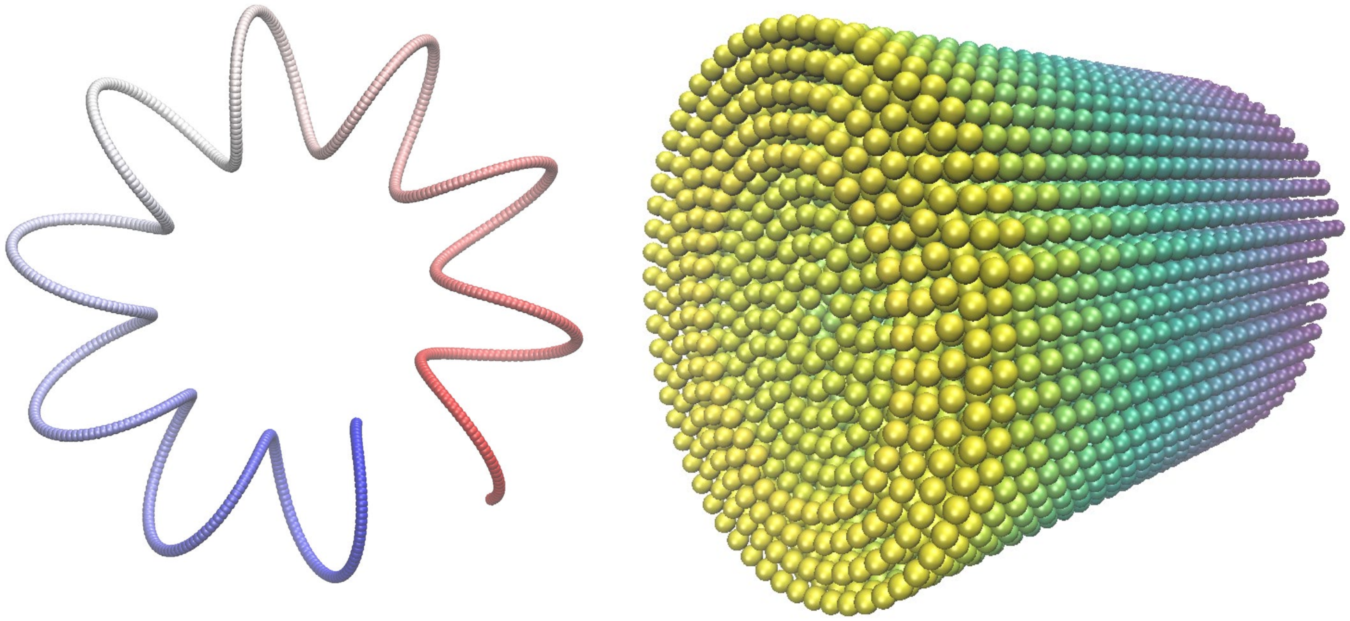

Figure S16 Initial configurations for simulations.

Left: Initial spiral spring conformation for simulations of 740-bead DT40 Chromosome 7.

Right: Initial spiral structure for simulations of “extended” DT40 mitotic chromosome 7 consisting of 10 copies of chromosome 7 concatenated head-to-tail.
